# The ERCC6L2-MRI-KU complex coordinates NHEJ at staggered DNA double-strand breaks

**DOI:** 10.1101/2025.11.28.691009

**Authors:** Pia I. Reichl, Ceylan Sonmez, Yueru Sun, Ashleigh King, Jean S. Metson, Benjamin Davies, Adam C. Wilkinson, Francisca Lottersberger, J. Ross Chapman

## Abstract

ERCC6L2 disease is a recessive bone marrow failure (BMF) syndrome caused by mutations in the SNF2-like putative DNA helicase ERCC6L2. While implicated in DNA replication, double strand break (DSB) repair via non-homologous end joining (NHEJ), and interstrand crosslink (ICL) repair, how ERCC6L2 supports haematopoietic longevity remains unclear. Investigating this *in vivo*, we find that an *Ercc6l2-*deficient haematopoietic stem and progenitor cell (HSPC) compartment in mice is unexpectedly resilient. *Ercc6l2* loss was also tolerated in mice co-deficient for endogenous formaldehyde detoxification, which precipitates early-onset BMF in models of Fanconi anaemia. Instead, *Ercc6l2*-deficient mice display a mild immunodeficiency, arising from defects in immunoglobulin class-switch recombination (CSR), that synergise with shieldin-deficiency, implicating ERCC6L2 and shieldin in distinct repair mechanisms. Furthermore, we demonstrate that ERRC6L2 stimulates chromosome fusions in the context of staggered, but not blunt dysfunctional telomeres. We reconcile ERCC6L2’s NHEJ function through proteomic elucidation of its endogenous interactome and AlphaFold structural modelling to reveal a complex formed of ERCC6L2 and KU that is bridged by the NHEJ accessory factor MRI/CYREN. Consequently, ERCC6L2-MRI inter-dependence characterises CSR. Together, our findings implicate the ERCC6L2-MRI complex as a KU-regulatory DNA translocase coordinating classical-NHEJ at staggered-end DSBs. We suggest that similar staggered-end breaks represent the pathological substrates driving haematopoietic failure in ERCC6L2 disease.

## INTRODUCTION

Inherited bone marrow failure syndromes are a heterogeneous group of disorders characterised by bone marrow failure (BMF) and typically one or more other somatic abnormalities, as well as a predisposition to developing cancer^1^.They are usually grouped into three classes based on the underlying cellular defect causing the failure, namely ribosome biogenesis, telomere maintenance, or DNA repair, such as in case of Fanconi anaemia.

Recently, inherited mutations affecting the gene encoding the SNF2-like putative helicase ERCC6L2 were shown to result in a BMF syndrome coined ERCC6L2 disease^2,3^. Clinical features of ERCC6L2 disease include mild changes in peripheral blood counts, yet severe bone marrow hypoplasia, frequent somatic *TP53* mutations and a high likelihood of disease progression to myelodysplastic syndrome (MDS) and acute myeloid leukaemia (AML), most commonly of the erythroid lineage^4,5^.

At a mechanistic level, the ERCC6L2 translocase has been implicated within multiple DNA repair pathways, including transcription-coupled nucleotide excision repair (TC-NER)^6^, ICL repair^3^, and DNA double-strand break (DSB) repair^7–9^. In a separate study, ERCC6L2 was also shown to assist DNA replication through centromeric DNA^10^. Most recently, unbiased genome-wide CRISPR-Cas9 screening approaches linked ERCC6L2 to DSB repair as an accessory factor in nonhomologous end joining (NHEJ)^7–9^. However, the mechanism through which ERCC6L2 assists NHEJ, and the contexts in which its activities become important remain ill-defined. Furthermore, how and why losses in its function specifically manifest in failed haematopoiesis and BMF in humans remains similarly unclear.

To address these questions, we generated and characterised the effect of ERCC6L2-deficency *in vivo* in mice. Our findings demonstrate that haematopoiesis remains surprisingly resilient in *Ercc6l2* knockout mice even in a sensitized background deficient in endogenous formaldehyde detoxification that precipitates HSPC attrition and BMF in mouse models of Fanconi anaemia^11^. Dispelling the notion of ICL repair defects as being causative of BMF in ERCC6L2 disease, we turned our focus to its function in NHEJ. Here, we show *Ercc6l2*-deficient mice present with mild immunodeficiency due to a context-specific NHEJ defect that specifically affect class switch recombination (CSR) but leaves V(D)J recombination intact. Using genetic epistasis to investigate where ERCC6L2 functions, we reveal it acts downstream of 53BP1, promoting the joining of AID-dependent DSBs by classical-NHEJ (c-NHEJ), in parallel to the shieldin-CST-Pol⍺-Primase DSB repair axis. Combining proteomics, structural-modelling and biochemical approaches, we reveal ERCC6L2 acts as a NHEJ-regulatory enzyme that associates with the DSB end-bound KU heterodimer, via bridging interactions provided by the NHEJ accessory factor MRI (CYREN). Lastly, we report experiments that reveal the context-specificity of ERCC6L2-participation in c-NHEJ, by revealing it stimulates the fusion of dysfunctional telomeres when they have G-strand ssDNA tailed, but not blunt telomeric ends.

Taken together, our findings implicate ERCC6L2-MRI as accessory translocase assisting KU-dependent DNA synapsis, and downstream joining at staggered-end DSBs. Our results thus reconcile its physiological importance at AID-dependent DSBs during CSR and implicate defects in the coordination of KU at endogenous ssDNA-tailed DSBs as the likely driving force of HSC attrition and dysfunction in ERCC6L2 disease.

## RESULTS

### Resilient haematopoiesis characterises a mouse model of ERCC6L2 disease

To better understand the role of ERCC6L2 in haematopoiesis, mice deleted for critical exons 2-16 of *Ercc6l2*, which compromises the ATP-binding and ATPase domain, were generated in an inbred C57BL/6J background (Fig. 1a and Extended Data Fig. 1a). Genomic DNA analysis first identified F0 founder mice harbouring the desired *Ercc6l2^em^*^1^ deletion allele, with reduced and ablated *Ercc6l2* mRNA expression reconfirmed in heterozygous and homozygous descendants of their backcrossed and intercrossed progeny, respectively (Extended Data Fig. 1b). *Ercc6l2^-/-^* mice were born at expected Mendelian frequencies and showed no overt phenotype, with no evidence of haematopoietic stem and progenitor cell (HS(P)C) attrition within the bone marrow of a small cohort of *Ercc6l2^-/-^* mice aged to 2 years (Extended Data Fig. 1c).

**Figure 1.**
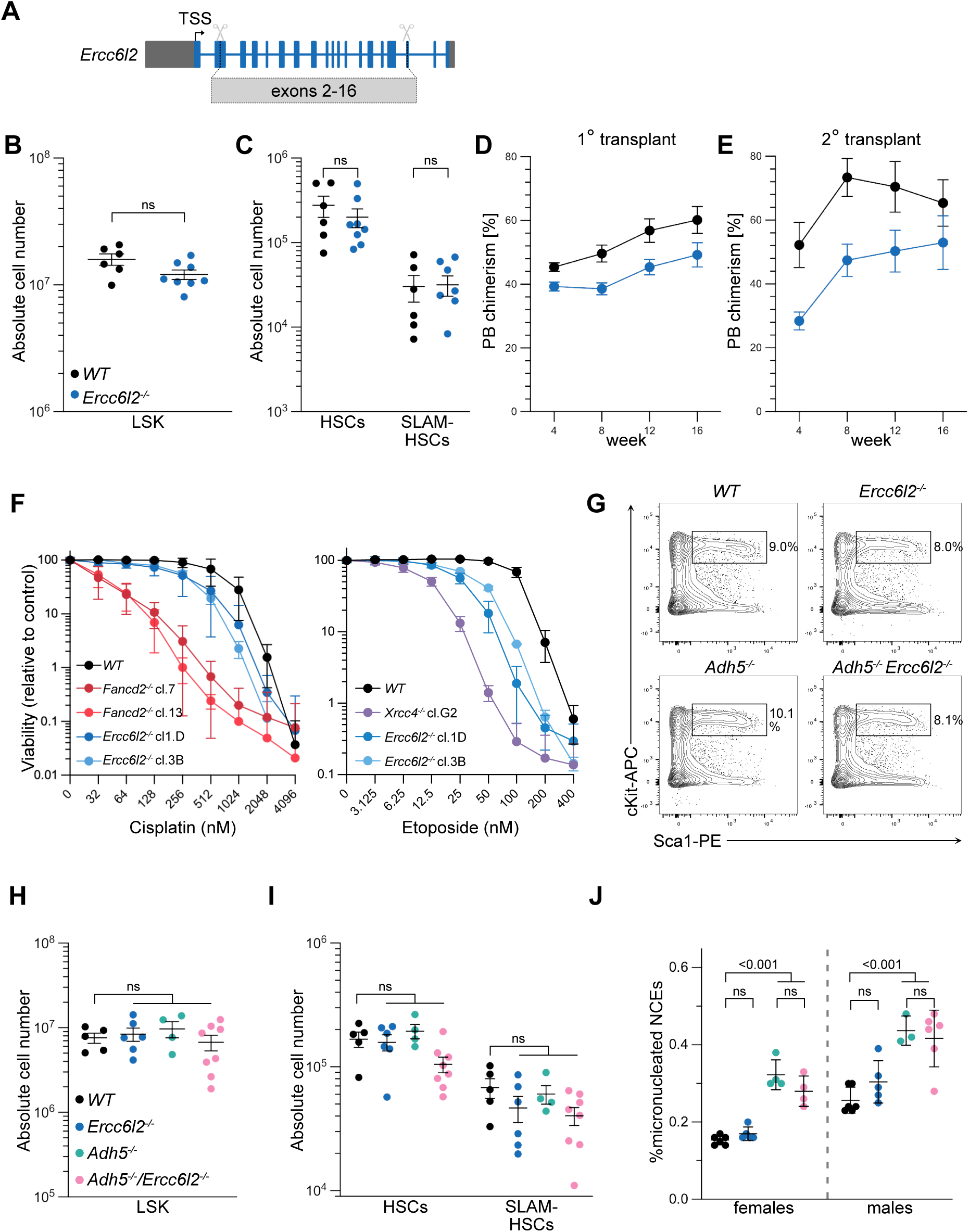
Resilient Haematopoiesis characterises a mouse model of ERCC6L2 disease. **A.** Schematic representation of *Ercc6l2* (knockout) allele generated for this study. TSS, transcription start site; grey, non-coding exons; blue, coding exons **B.** Absolute number of Lin^-^ Sca1^+^ c-Kit^+^ (LSK) cells in the bone marrow (one femur and one tibia) (*n*= 6-8 mice per genotype, where each data point is a single mouse). Significance was determined by an unpaired two-tailed *t*-test (mean ± SEM). ns, not significant **C.** Absolute number of CD34^-^ LSK cells (HSCs) and CD34^-^ CD48^-^ CD150^+^ LSKs (SLAM-HSCs) in the bone marrow (one femur and one tibia) (*n*= 6-8 mice per genotype, where each data point is a single mouse). Significance was determined by an unpaired two-tailed *t*-test (mean ± SEM) **D.** Primary competitive bone marrow transplantation assay was performed by transplanting 5 x 10^5^ wild-type or *Ercc6l2^-/-^* whole bone marrow cells (CD45.2) together with 5 x 10^5^ competitor cells (CD45.1) into irradiated recipients (CD45.1/CD45.2). Plots show donor vs. competitor chimerism in mononucleated peripheral blood (PB) (*n*= 4-5 mice per test condition, mean ± SEM). **E.** Secondary competitive bone marrow transplantation assay was performed by transplanting 1 x 10^6^ whole bone marrow cells from primary transplant recipients into irradiated recipients (CD45.1/CD45.2). Plots show donor vs. competitor chimerism in mononucleated peripheral blood (PB) (*n*= 4-5 mice per test condition, mean ± SEM). **F.** Survival of CH12-F3 cells grown for 5 days in the presence of indicated doses of cisplatin or etoposide (*n*= 3 biological replicates, with 3 technical replicates each, mean ± SD). **G.** Representative flow cytometry plots showing Lin^-^ cells used to quantify LSK frequencies in the bone marrow. **H.** Absolute number of Lin^-^ Sca1^+^ c-Kit1^+^ (LSK) cells in the bone marrow (one femur and one tibia) (*n*= 5-8 mice per genotype, where each data point is a single mouse). Significance was determined by an unpaired two-tailed *t*-test (mean ± SEM). ns, not significant **I.** Absolute number of CD34^-^ LSK cells (HSCs) and CD34^-^ CD48^-^ CD150^+^ LSKs (SLAM-HSCs) in the bone marrow (one femur and one tibia) (*n*= 5-8 mice per genotype, where each data point is a single mouse). Significance was determined by an unpaired two-tailed *t*-test (mean ± SEM) **J.** Percentage of micronucleated normochromatic erythrocytes (NCEs) in the peripheral blood. Blood was taken from 8-12 week old mice via tail vein (n= 3-6 mice of each sex per genotype, where each data point is a single mouse). Significance was determined by an unpaired two-tailed *t*-test (mean ± SEM).

Given its dysfunction in human ERCC6L2 disease, we focussed our initial investigations on the haematopoietic stem and progenitor compartment of *Ercc6l2^-/-^* mice, analysing these for evidence of stem cell defects or attrition. Flow cytometric quantification of haematopoietic stem and progenitor cells in *Ercc6l2^-/-^*mice revealed wild type frequencies of the Lin^-^ Sca1^+^ c-Kit^+^ (LSK) cell population in the bone marrow (Fig. 1b). Similarly, enumeration of immunophenotypic HSCs (CD34^-^ LSKs) and long-term SLAM-HSCs (CD48^-^ CD150^+^ LSKs) in *Ercc6l2^-/-^* mice revealed normal frequencies in both subsets (Fig. 1c). In consideration of sub-penetrant defects in HSC fitness or function in *Ercc6l2-*deficient mice, we performed competitive bone marrow transplantation experiments, in which equal mixes of *Ercc6l2^-/-^*(CD45.2) and wild-type (CD45.1/CD45.2) competitor whole bone marrow cells were injected intravenously into lethally irradiated CD45.1 wild-type recipient mice. Here, transplanted *Ercc6l2^-/-^* bone marrow successfully engrafted and repopulated the bone marrow of irradiated recipient mice, where it sustained peripheral blood production at levels equivalent to that of wild-type competitor BM in both primary (Fig. 1d), and secondary (serial) transplant experiments (Fig. 1e). In all cases, no differences between the peripheral blood lineage outputs of *Ercc6l2^-/-^* or wild type mice were evident (Extended Data Fig. 1d-g), and their stem cell engraftment efficiency was comparable in both cases (Extended Data Fig. 1h, i). Taken together, these results suggest that in mice, ERCC6L2 is largely dispensable for HSC self-renewal and long-term reconstitution of haematopoiesis.

The preserved haematopoietic capacity of *Ercc6l2*-knockout mice suggested that drivers of HSC attrition in human ERCC6L2 disease patients may be absent or attenuated in mice. A similar species divergence is observed in Fanconi anaemia (FA) where, despite equivalent defects in ICL repair, the phenotypes of aplastic anaemia, BMF, and haematological malignancies are sub-penetrant in murine models compared to patients^12^. Interestingly, ERCC6L2-deficient patient fibroblasts were shown to be hypersensitive to treatments with the ICL-inducing agent mitomycin C, albeit to a lesser degree than FA patient fibroblasts^3^. Indeed, ICL repair defects are suggested in *Ercc6l2*-knockout clones of the mouse B lymphoblastoid cell-line CH12-F3, which exhibited intermediate cisplatin sensitivity relative to *Fancd2-*knockout CH12-F3 clones (Fig. 1f), in addition to a sensitivity to the Topoisomerase 1 inhibitor etoposide, which suggested defects also in DSB repair (Fig. 1f and Extended Data Fig. 2a).

This sensitivity of *Ercc6l2-*deficient mouse cells to ICL-inducing treatments prompted us to first consider the specific contribution of mild ICL-repair defects to the progressive HSC attrition in *ERCC6L2-*deficient patients, where BMF is less severe and follows a much longer latency than in FA^4,13^. To test this, we interbred *Ercc6l2* knockout mice with mice harbouring knockout alleles for the alcohol dehydrogenase 5 (*Adh5*) gene^14^, which encodes the primary mammalian enzyme mediating the detoxification and clearance of endogenous formaldehyde (Extended Data Fig. 2b)^15^. In *Adh5*-deficient mice, increased intracellular formaldehyde accumulation drives DNA damage to levels that synergise severely with ICL-repair defects in *Fancd2-*deficient mice^11^. As a consequence, *Fancd2^-/-^ Adh5^-/-^* mice typically die within 3-6 weeks of birth due to aplastic anaemia and bone marrow failure, precipitated by unrepaired formaldehyde-induced DNA damage driving p53-dependent ablation of HSPC cell pools^11,16^. However, when we derived *Adh5^-/-^ Ercc6l2^-/-^* mice, they were born showing no signs of disrupted haematopoiesis. The frequencies of HSPCs and HSCs in 8-12 week-old *Adh5^-/-^Ercc6l2^-/-^* mice were comparable to that in single knockout control mice (Fig. 1g-i and Extended Data Fig. 2c-d). Furthermore, when levels of micronucleated normochromatic erythrocytes (MN+ NCEs) were used as a proxy of genomic instability in the peripheral blood in these strains^17^, although elevated versus wild type, they occured at equivalent MN+ NCE levels in *Ercc6l2^-/-^ Adh5^-/-^* and *Adh5^-/-^* controls (Fig. 1j and Extended Data Fig. 2e). As such, we observed no evidence of synergy between formaldehyde-induced DNA damage in *Ercc6l2-*deficient mice, a result markedly different to *Fancd2^-/-^ Adh5^-/-^* mice^18^. Thus, while formaldehyde-induced interstrand crosslinks drive bone marrow failure in mouse models of FA, we surmise that aldehyde-driven ICLs were unlikely to contribute to HSPC attrition in ERCC6L2 disease. These findings shifted our focus toward the DSB repair functions of ERCC6L2 as a more plausible determinant of disease pathology.

#### ERCC6L2 promotes classical-NHEJ at AID-dependent DSBs in CSR

In developing and stimulated lymphocytes NHEJ supports the generation of functional antigen receptor genes through two distinct mechanisms: V(D)J recombination – through which functional antigen receptors are assembled from germline *variable*, *diversity* and *joining* gene segments by deletional NHEJ in developing B and T lymphocytes; and immunoglobulin (Ig) class-switch recombination (CSR) – through which antibody isotype-encoding constant (C) gene segments are excised and replaced within the immunoglobulin heavy chain locus (*Igh*) in peripheral mature B cells^19^. Consistent with ERCC6L2’s attributed role in NHEJ^7–9^, Ig class-switching defects characterised a recently published *Ercc6l2-*knockout mouse, where they reportedly coincided with non-productive inversional recombination events at the *Igh*^7^. Indeed, an equivalent magnitude of defect was faithfully replicated when we measured CSR in stimulated B splenocytes from our *Ercc6l2^-/-^* knockout mice, with intermediate CSR reductions also apparent in B cells from heterozygous animals (Extended Data Fig. 3a).

ERCC6L2’s involvement in CSR could implicate it in the 53BP1 pathway, which orchestrates specialized DNA end-processing and ligation activities essential for joining DSBs induced by activation-induced deaminase (AID). Extensive nucleolytic processing of AID-dependent clustered U:G mismatches generate staggered cleavages at the *Igh* locus, producing 5′-recessed single-stranded DNA termini that are inherently incompatible with direct NHEJ^20^ . Productive joining therefore relies predominantly on 53BP1-dependent promotion of fill-in synthesis, mediated through recruitment of shieldin and the CTC1-STN1-TEN1 (CST) complex, which direct POLA-PRIM1-dependent priming and processive DNA synthesis involving polymerases such as POLζ^8,21–25^. Hypothesizing an auxiliary role for ERCC6L2 helicase and/or translocase activity in the stimulation of these processing steps, we interbred mice to generate strains in which *Ercc6l2* was co-deleted for either *53bp1*, or *Shld*2, which encodes the core DNA binding subunit of shieldin (Extended Data Fig. 3b). Resulting *Ercc6l2^-/-^ 53bp1^-/-^*and *Ercc6l2^-/-^ Shld2^-/-^* mice were viable, born at expected frequencies, and again presented with no overt phenotypes. Moreover, mice co-deficient in *Ercc6l2* and *53bp1*, showed only a marginal decrease in the total number of B cells in the bone marrow (Fig. 2a), and no decrease in the total number of B cells in the spleen when compared to animals lacking *53bp1* alone (Fig. 2b). Marginal decreases in bone marrow B cells were not associated with decreases at any particular developmental stage (Extended Data Fig. 3c-f), suggesting that ERCC6L2 does not participate to V(D)J recombination in the absence of 53BP1. This assumption was also supported by the behaviour of thymic T cells, where the total number of thymocytes was also slightly decreased (Extended Data Fig. 4a), but without defects at any particular developmental stage (Extended Data Fig. 4b-c). ERCC6L2’s participation in V(D)J recombination was formally ruled out when we examined RAG-induced V(D)J recombination frequencies directly with the pINV-GFP reporter substrate^13^ in *v-abl* kinase transformed pre-B cell lines generated from the bone marrow of *Ercc6l2^-/^*^-^ and *Ercc6l2^-/-^ 53bp1^-/-^* mice (Extended Data Fig. 4d-e). Similarly, *Ercc6l2^-/-^ Shld2*^-/-^ mice showed normal cell frequencies across all stages of B cell development in the bone marrow (Fig. 2c and Extended Data Fig. 5a, c) and B cell maturation in the spleen (Fig. 2d and Extended Data Fig. 5b, d).

**Figure 2.**
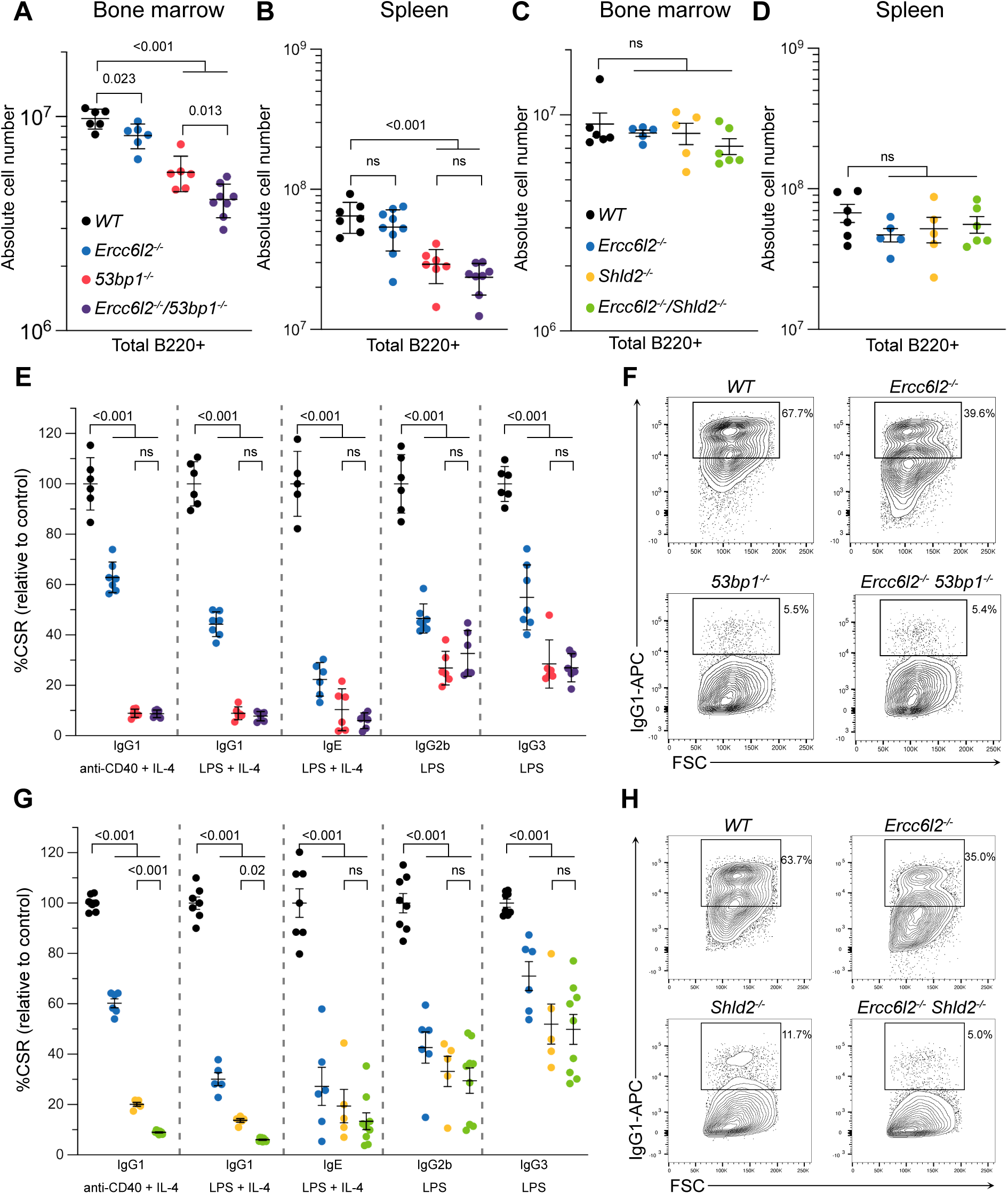
Synergistic non-redundant NHEJ activities of ERCC6L2 and shieldin support antibody class-switching. **A.** and **C.** Absolute number of B220^+^ B cells in the bone marrow (one femur and one tibia (*n*= 5-8 mice per genotype, where each data point is a single mouse). Significance was determined by an unpaired two-tailed *t*-test (mean ± SEM). ns, not significant **B**. and **D**. Absolute number of B220^+^ B cells in the spleen (*n*= 5-8 mice per genotype, where each data point is a single mouse). Significance was determined by an unpaired two-tailed *t*-test (mean ± SEM). ns, not significant **E.** and **G.** Splenic B cells cultured with indicated cytokines for 96 h and stained for IgG1, IgE, IgG2b, or IgG3. (*n*= 5-9 mice per genotype, where each data point represents a single mouse). Isotype switching frequency was normalized to two wild-type mice for each experiment. Significance was determined by a one-way ANOVA with Tukey’s correction (mean ± SEM). **F.** and **H.** Representative flow cytometry plots showing splenic B cells cultured with anti-CD40 and IL-4 for 96 h and stained for IgG1.

The development and maturation of B cells in *Ercc6l2^-/-^ 53bp1^-/-^* and *Ercc6l2^-/-^ Shld2^-/-^*mice and their single knockout and wild type controls (Fig. 2a-d and Extended Data Fig. 3, 5), allowed us to focus on ERCC6L2’s functional interplay with 53BP1 and shieldin during CSR. As expected^26,27^, the induction of CSR – across all Ig isotypes – was near-completely ablated in *53bp1^-/-^* B splenocytes, with no additive effect in *Ercc6l2^-/-^ 53bp1^-/-^* B cells (Fig. 2e-f and Extended Data Fig. 6a-c). However, the absence of apparent synergistic CSR defects in *Ercc6l2^-/-^ 53bp1^-/-^* B cells could not be interpreted confidently as epistasis given the severity of CSR defects in *53bp1^-/-^* control mice. By comparison, B cells from shieldin-deficient mice harboured milder defects with significant residual class-switching to IgG1 (Fig. 2g-h, Extended Data Fig. 7a). Comparing CSR frequencies in stimulated splenic B cells from *Shld2^-/-^, a*nd *Shld2^-/-^ Ercc6l2^-/-^* mice, showed combined loss of *Ercc6l2* and *Shld2* caused a total collapse of CSR to IgG1 (Fig. 2g-h) - equivalent to loss of *53bp1* - that was neither accompanied by cell proliferation nor cell survival defects (Extended Data Fig. 7b-c). Thus, our findings indicate that ERCC6L2 and the shieldin complex mediate independent, non-redundant joining activities at AID-dependent DSBs.

#### ERCC6L2 promotes fusions at staggered, but not blunt, telomeric DSBs

As AID-dependent DSBs have ends with heterogenous overhang, the specific importance of ERCC6L2 for some, shieldin independent, DNA end-joining events during CSR led us to consider whether ERCC6L2 supports NHEJ specifically in the context of DSBs with recessed termini. To this end, we investigated ERCC6L2’s potential role in LIG4-dependent c-NHEJ of dysfunctional telomeres with different end structures as model substrates. In mammals, telomeres are protected from being recognized as DSBs by the binding of the 6-protein shelterin complex onto an array of TTAGGG repeats followed by a 50-300 nt long 3’ overhang that folds back to form a lariat-like structure called a t-loop^28^. The shelterin protein TRF2 is essential for t-loop formation and telomere protection^29–31^. In its absence, the telomeres, which retain their 3’ overhangs, elicit a DSB-dependent ATM response and are engaged by c-NHEJ, resulting in the widespread accumulation of intrachromosomal fusions^29–31^. In addition, TRF2 recruits the shelterin component RAP1 and the exonuclease APOLLO to restrain DNA-PK and promote the 3’ overhang formation, respectively, at the blunt telomeres generated by the leading strand DNA replication^30,32–39^. While APOLLO-deficient blunt leading end telomeres are fused by alt-EJ in S/G2, resulting in a modest increase of chromatid fusions^36–38,40^, the additional deletion of RAP1 causes a significant increase in LIG4-dependent intrachromosomal fusions^39^. Therefore, to formally address ERCC6L2’s contribution to c-NHEJ at blunt versus recessed ssDNA-comprising termini, we investigated its contribution to the fusion of dysfunctional telomeres with or without overhang due to the deletion of, respectively, *Trf2* (Fig. 3a, top) or both *Rap*1 and *Apollo* (Fig. 3a, bottom). In agreement with previous findings^30,39^, telomere fluorescence in situ hybridization (FISH) in the control MEFs after the deletion of *Trf2* by CRISPR/Cas9 showed that most telomeres were engaged in chromosome-type fusions (Fig. 3b-d and Extended Data Fig. 8a), while the deletion of both *Rap1* and *Apollo* caused chromosome fusions at about 25% of telomeres in addition to the about 2% chromatid fusions due to Apollo sole deletion (Fig. 3e-g and Extended Data Fig. 8b-d). In *Ercc6l2-*deficient MEFs there was a significant reduction in the number of chromosome fusions after *Trf2* deletion (Fig. 3b-d), but not after the combined *Rap1*/*Apollo* deletion (Fig. 3e-g). Furthermore, *Ercc6l2* deletion status was not found to influence chromatid fusion frequencies when *Apollo* was deleted in isolation (Extended Data Fig. 8b-c), excluding a significant role for ERCC6L2 in the APOLLO-dependent processing of newly replicated leading-end telomeres, the protection of the same ends from MRN-dependent resection, or the repair by alt-EJ^40–42^.

**Figure 3.**
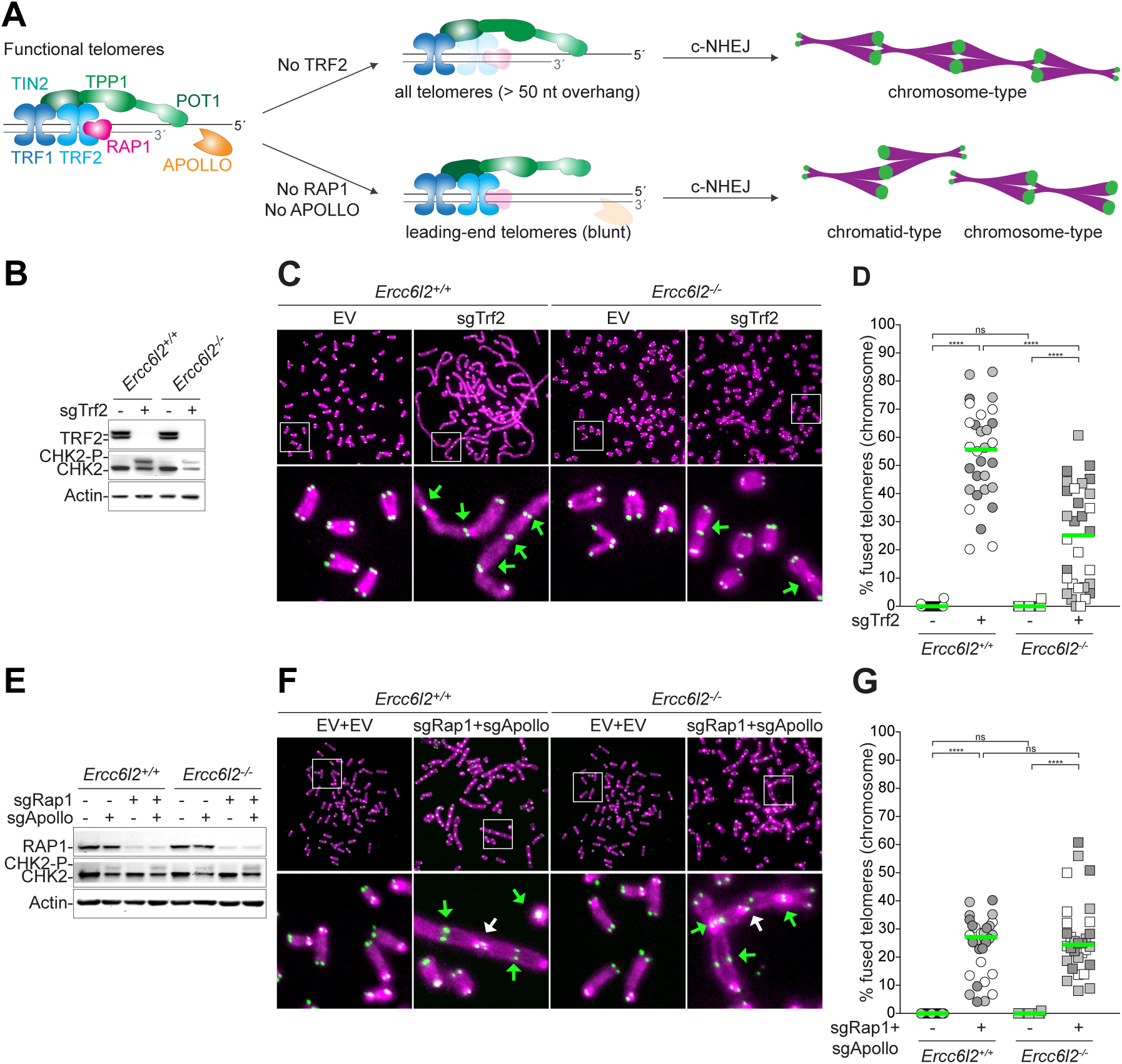
ERCC6L2 stimulates c-NHEJ at staggered, but not blunt dysfunctional telomeres. **A.** c-NHEJ at dysfunctional telomeres with and without overhang. Functional telomeres are formed by a repetitive array of TTAGGG repeats ending with a long (50-300 nt) 3’ overhang bound by the shelterin complex (TRF1, TRF2, RAP1, TIN2, TTP1, POT1). At newly replicated leading end telomeres the overhang is formed by the Exonuclease APOLLO. In the absence of TRF2, telomeres still retaining their overhang are fused by c-NHEJ (top). In the absence of APOLLO and RAP1, the blunt telomeres resulting from replication cannot be resected and are fused by c-NHEJ (bottom). **B.** Immunoblot for TRF2 deletion, CHK2 phosphorylation as indication of ATM activation and actin as loading control in *Ercc6l2^+/+^* and *Ercc6l2^-/-^* MEFs 108 h after transduction with single guide RNA (sgRNA) targeting Trf2 (sgTrf2). **C.** Representative FISH of metaphase spreads of *Ercc6l2* positive and negative 108 h after *Trf2* deletion. Telomeres were detected with Cy3-(TTAGGG)3 (green) and DNA was stained with DAPI (magenta). Green arrows highlight chromosome-type fusions. **D.** Quantification of telomeres involved in chromosome fusions per metaphase after deletion of *Trf2* as in (B-C). Data from 3 independent experiments, 10 metaphases per experiment (n = 30 total), with median. **E.** Immunoblot for *Rap1* deletion, CHK2 and actin in *Ercc6l2^+/+^* and *Ercc6l2^-/-^* MEFs 120 h after transduction with single guide RNA (sgRNA) targeting Rap1 (sgRAp1) and/or Apollo (sgApollo). **F.** Representative FISH of metaphase spreads of *Ercc6l2* positive and negative MEFs 120 h after *Apollo* and *Rap1* deletion. Telomeres were detected with Cy3-(TTAGGG)3 (green) and DNA was stained with DAPI (magenta). Green arrows highlight chromosome-type fusions. White arrows highlight chromatid-type fusions. **G.** Quantification of telomeres involved in chromosome fusions per metaphase after deletion of *Rap1* and *Apollo* as in (e-f). Data from 2 (no sgRNAs) or 3 (sgRap1 + sgApollo) independent experiments, 10 metaphases per experiment with median. Ordinary one-way analysis of variance (ANOVA). *P ≤ 0.05, **P ≤ 0.01, ***P ≤ 0.001, ****P ≤ 0.0001; ns, not significant.

Overall, these data support our first prediction regarding ERCC6L2’s specific ability to stimulate c-NHEJ in the context of staggered DNA ends, pointing towards a role in assisting DNA end-processing events at ligation-refractory ssDNA-tailed termini to enable KU/LIG4-dependent ligation and c-NHEJ.

#### Discovery and characterization of the ERCC6L2-MRI-KU complex

To interrogate the mechanism of ERCC6L2 regulated c-NHEJ at staggered DSBs, we used proteomics to define the endogenous ERCC6L2 interactome. To achieve this, we used CRISPR-Cas9 to modify both alleles of the *Ercc6l2* gene to encode N-terminal TwinStrep-HA-tags in CH12-F3 murine B cells (Extended Data Fig. 9a). Due to ERCC6L2’s extreme low expression, lysates prepared from cultures of ∼1×10^8^ *Ercc6l2^Strep-HA/Strep-HA^*cells were used to isolate ERCC6L2 complexes by affinity purification on Strep-TactinXT™ resin following X-ray cell irradiation or mock treatment. Biotin eluted complexes were then subjected to analysis by liquid chromatography-tandem mass spectrometry (LC-MS/MS) over two independent biological replicates (Fig. 4a), with successful purification confirmed by immunoblotting (Extended Data Fig. 9b). Besides the numerous ERCC6L2 peptides detectable by LC-MS/MS, the only other peptides reproducibly detected in Strep-HA-ERCC6L2 precipitates corresponded to the KU70 and KU80 subunits of DNA-PK, and the NHEJ accessory factor MRI/CYREN. These are previously reported interacters of ERCC6L2^7^, that we found to be co-purified in endogenous Strep-HA-ERCC6L2 complexes under both untreated and cell irradiation conditions (Table 1 and STable 1). The existence of a KU-ERCC6L2 complex could also also be validated by immunoblotting across smaller-scale batch Strep-Tactin purifications (Extended Data Fig. 9c).

**Figure 4.**
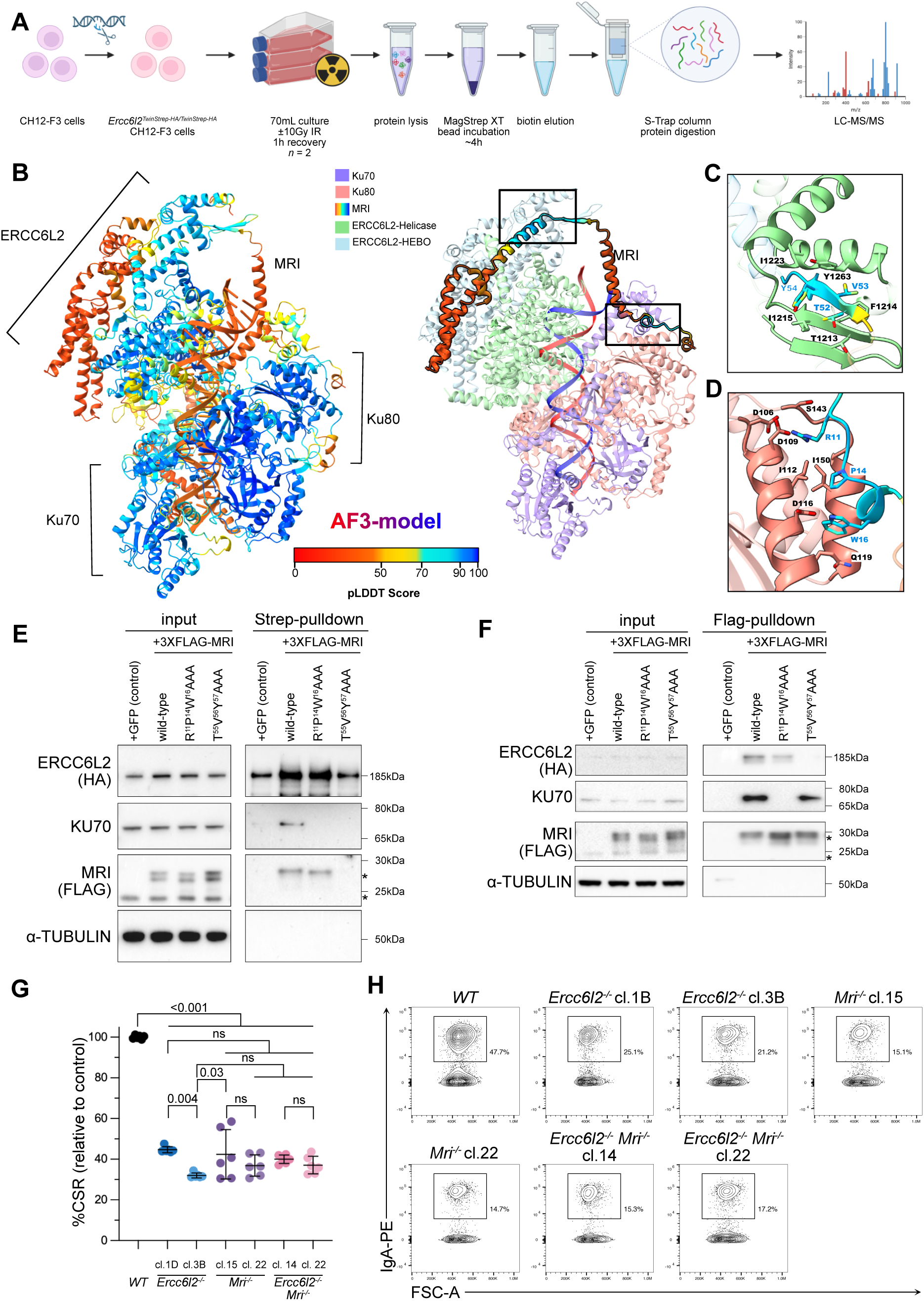
The ERCC6L2-MRI-KU complex supports classical NHEJ in CSR. **A.** Schematic representation of TwinStrep-HA-ERCC6L2 purification and proteomic analysis strategy. Briefly, the endogenous *Ercc6l2* locus was biallelically tagged with a TwinStrep-HA tag. 70 mL cell cultures were either mock-treated or irradiated (10 Gy) and lysates were collected following a 1h recovery. ERCC6L2 was purified using MagStrep XT beads and eluted in biotin. The elutions were then captured and tryptic-digested on S-Trap columns and the resulting peptides were analysed using LC-MS/MS. This schematic was generated using BioRender. **B.** Top ranking ERCC6L2-MRI-Ku-DNA complex predicted structure coloured according to plDDT score (left) and by individual subunits (right), in which MRI highlighted in plDDT colour, ERCC6L2 helicase domain in pale green, ERCC6L2-HEBO domain in pale blue, Ku70 in purple and Ku80 in coral. The potential MRI-ERCC6L2 interaction motif and MRI-Ku80 binding motif are highlighted. **C.** Zoomed-in ERCC6L2-MRI binding site, interacting residues on MRI and ERCC6L2 are labelled. **D.** Zoomed-in Ku80-MRI binding site with interface residues showed in stick and labelled, coloured in the same scheme as right panel in B. **E.** Western blot analysis of CH12-F3 lysates from TwinStrep-ERCC6L2 pulldown experiments. Blot is representative of *n*=2 biological replicates. **F.** Western blot analysis of CH12-F3 lysates from FLAG-MRI pulldown experiments. Blot is representative of *n*=2 biological replicates. **G.** IgM-to-IgA CSR frequencies in *Ercc6l2* and *Mri* knockout CH12-F3 cells normalized to wild-type cells (*n*=3 biological replicates, with two technical replicates per experiment). Significance was determined by a one-way ANOVA with Tukey’s correction (mean ±SD). **H.** Representative flow cytometry plots **H.** Representative flow cytometry plots for IgM-to-IgA CSR frequencies.

**Table 1.** List of antibodies for the biotin-conjugated lineage antibody stain. Working dilutions are listed.

| Antibody | Manufacturer | Catalogue number | Dilution |
| --- | --- | --- | --- |
| B220-biotin | invitrogen | 13-0452-85 | 1:1400 |
| CD127-biotin | invitrogen | 13-1271-85 | 1:1400 |
| CD4-biotin | invitrogen | 13-0042-85 | 1:2800 |
| CD8-biotin | invitrogen | 13-0081-86 | 1:2800 |
| Ly6G-biotin | invitrogen | 13-5931-85 | 1:700 |
| Ter119-biotin | invitrogen | 13-5921-85 | 1:700 |

**Table 1.** ERCC6L2 interactome detected via LC-MS/MS. Interacting proteins were determined as in Fig. 3a (*n*=2 independent experiments per genotype and condition). Peptide identifications were accepted if they could be established at >95.0% probability, and protein identifications were accepted if they could be established at >99.0% probability. Full table of LC-MS/MS results is available in the supplementary files.

| Protein symbol | Coverage (%) | Total unique spectra |  |  |  |  |  |  |  |
| --- | --- | --- | --- | --- | --- | --- | --- | --- | --- |
|  |  | R1_T<br>S-HA-ERCC<br>6L2 | R1_TS-HA-ERCC6<br>L2<br>+IR | R2_TS-HA-ERCC6<br>L2 | R2_TS-HA-ERCC6<br>L2<br>+IR | R1_<br>CTR | R1_<br>CTR<br>+IR | R2_<br>CTR | R2_<br>CTR<br>+IR |
| ERCC6<br>L2 | 13 | 14 | 21 | 12 | 16 | 0 | 0 | 0 | 0 |
| KU80<br>(Xrcc5) | 6.3 | 3 | 4 | 3 | 3 | 0 | 0 | 0 | 0 |
| KU70<br>(Xrcc6) | 5.9 | 3 | 4 | 3 | 1 | 0 | 0 | 0 | 0 |
| CYREN<br>(Mri) | 14 | 1 | 2 | 2 | 2 | 0 | 0 | 0 | 0 |

To study the molecular basis of ERCC6L2-MRI interaction with DNA-bound KU, we performed multiple rounds of structure modelling using AlphaFold3 (AF3)^43^. Strikingly, AF3 generated high-confidence models in which ERCC6L2 formed a complex with DNA bound KU70-KU80, linked by bridging interactions provided by ERCC6L2- and KU- binding interfaces in MRI. Superimposition of all the structural models enabled us to evaluate the prediction variability at these interfaces. The root means square deviation (r.m.s.d.) at ERCC6L2-MRI interface was 0.708±0.14Å, indicating the regions are highly consistent among all predicted models. Together with the low expected prediction error (PAE) score (<15) and high pLDDT (>90) (Fig S9d, e), this indicates that the interface prediction is likely accurate. Of special interest in the predicted structure were high-confidence interactions between ERCC6L2-MRI, and MRI-KU. Specifically, a β-strand located within central sequences in MRI binds to the C-terminal HEBO domain in ERCC6L2 by forming an intermolecular β-sheet with two β-strands in this domain (Fig. 4b). The antiparallel ß-sheet is stabilised by a backbone hydrogen-bonding network formed between the corresponding ß-stands of MRI and ERCC6L2. The sidechains T52, V53 and Y54 on MRI pack against T1213, F1214 and I1215 on ERCC6L2 via hydrophobic contacts and aromatic ν-n stacking interactions. I1223 and Y1263 of ERCC6L2 buttress towards the hydrophobic core of the interface, reinforcing the inter-stand interaction (Fig. 4c).

AF3 models of the DNA bound ERCC6L2-MRI-KU complex also highlighted a previously characterised KU-MRI interaction, mediated by the Ku-Binding Motif (KBM) spanning R11, P14, and W16 in MRI^44,45^, and the von Willebrand domain (vWD) in Ku80^46^ In our model, the conserved KBM loop (MRI 11-20) is closely tethered to two α-helices of Ku80 vWD by several hydrogen bonds and salt bridges formed between D106, D109, D116, Q119, and S143 on Ku80, and R11 and W16 on MRI. Hydrophobic residues I112, I150 of Ku80 engage with MRI P40, further stabilising the interface (Fig. 4d).

To test the predicted interactions and investigate their interdependency, we generated *Mri^-/-^ Ercc6l2^Strep-HA/Strep-HA^* CH12-F3 cells, which were then virally reconstituted with triple-FLAG MRI fusion variants encoding alanine substitutions at the predicted Ku80 or ERCC6L2 HEBO domain contacting residues. Strep-HA pulldowns performed with lysates prepared from these cells showed that the ERCC6L2-KU interaction was strictly MRI-dependent, with KU absent from Strep-ERCC6L2 purifications from lysates prepared from MRI-deleted cells (Fig. 4e). Additionally consistent with the structural model, mutations in the KBM loop in MRI specifically abolished the interaction with KU, without impacting the MRI-ERCC6L2 interaction (Fig. 4e). Likewise, mutations in the predicted HEBO-interacting β-sheet in MRI abolished interactions between ERCC6L2 and MRI-KU, indicating that MRI acts to link KU and ERCC6L2, with Flag-MRI pulldowns revalidating the interactions defined in ERCC6L2 purifications (Fig. 4f).

Together, these structural and biochemical findings show MRI acts as a molecular bridge linking ERCC6L2 to KU on DNA, via conserved a HEBO-interacting β-strands and Ku80-binding motifs in MRI, respectively. These findings reveal an unexpected ERCC6L2-MRI interplay, that likely reconciles the roles of both proteins in NHEJ.

Given that CSR frequencies are reportedly reduced in *Mri-*knockout mice^47^, we next sought to define the functional relevance of MRI-ERCC6L2 interplay during this process^47–49^. To this end, we stimulated *Ercc6l2^-/-^*, *Mri^-/-^*, and *Ercc6l2^-/-^ Mri^-/-^* CH12-F3 cells, and assayed IgM to IgA class-switching relative to control cells. Indeed, class switching rates were equivalent in CH12-F3 cell lines lacking *Mri* and *Ercc6l2,* with no further additive defect detected in *Ercc6l2^-/-^ Mri^-/-^* cells (Fig. 4g-h). This indicates MRI-ERCC6L2 cooperativity underpins the function of both factors during NHEJ, establishing the role of the ERCC6L2-MRI-KU complex in the repair of subsets of AID-dependent DSBs during CSR.

Taken together, our findings allow us to make a coherent mechanistic prediction for ERCC6L2 function in c-NHEJ: anchored to KU by MRI at ligation-refractory ssDNA-tailed termini, the ERCC6L2 translocase likely repositions the c-NHEJ machinery to stimulate DNA end processing activities that licence efficient KU/LIG4-dependent ligation.

## DISCUSSION

In humans, *ERCC6L2* loss-of-function manifests in bone marrow failure, implicating ERCC6L2’s importance in the maintenance of HSCs that typically sustain lifelong blood production. Here, we make the paradoxical observation that in mice, an *Ercc6l2*-deficent HSPC compartment is unexpectedly resilient. Specifically, *Ercc6l2^-/-^* mice comprise normal populations of HSCs and HSPCs that exhibit a normal capability to sustain haematopoiesis under basal conditions, and that can engraft and repopulate functional bone marrow of lethally irradiated recipient mice over serial transplants. Remakably, HS(P)Cs even remain hardy in the face of chronic formaldehyde-induced genotoxic stress. In this scenario, the fact that aldehyde-clearance defects did not precipitate a haematopoietic phenotype in *Ercc6l2/Adh5* double-knockout mice clearly distinguishes ERCC6L2 disease from FA, where defective aldehyde-detoxification precipitates severe BMF through the rapid attrition of HSC pools perinatally. Thus, our findings implicate distinct endogenous mechanisms of HSC attrition in FA and ERCC6L2 disease causation.

Through the genetic dissection of the mild immunological phenotypes of *Ercc6l2^-/-^*mice, our findings, however, reveal an important function for ERCC6L2 during NHEJ. We also provide mechanistic insights into its basis that help reconcile why ERCC6L2 regulates NHEJ in the context of AID-dependent DSBs during CSR, but not in the case of RAG-induced DSBs during V(D)J recombination. More specifically, our findings suggest this can be explained by the different structural properties of DNA termini generate during each process.

During V(D)J recombination, the KU-DNA-PKcs holoenzyme engages the hairpin-sealed DSB intermediates of RAG1/2-induced cleavage, together with the XRCC4/XLF/PAXX c-NHEJ scaffolds. These assemblies form the core of the c-NHEJ synapsis complexes that can autonomously couple DNA end-tethering and alignment activities to all the necessary processing steps (i.e. hairpin-opening, DNA end polymerisation and cleanup) required to convert incompatible ends to those amenable to LIG4-dependent ligation^19^ . During CSR, by contrast, the fact that AID-induced DSBs arise following the nucleolytic excision and processing of interspaced uracil residues distributed between both strands within *Igh* S-repeat elements means that DSBs will ultimately possess ssDNA overhangs of various lengths and polarity. Accordingly, these DSBs necessitate specialised NHEJ accessory factors such as ERCC6L2. Our proposition that the presence of an overhang determines the requirement for ERCC6L2 during NHEJ is further substantiated by concordant observations at deprotected telomeres, where ERCC6L2 enhances fusion efficiency at TRF2-depleted 3’ ssDNA overhangs, yet offers no stimulation of NHEJ at blunt-ends. However, since our findings reveal that shieldin/CST and ERCC6L2 constitute synergistic, yet non-redundant DNA end joining mechanisms in the context of AID-dependent DSBs, ERCC6L2 could be implicated in the stimulation of c-NHEJ specifically in the context of staggered DSBs with short overhangs that either do not, or no longer, require extensive fill-in by shieldin-CST complexes. Altogether, the synergy between shieldin-CST and ERCC6L2 may provide the mechanistic versatility needed to prime ligation within overhangs of different length and likely overhang polarities (Fig. 5).

**Figure 5.**
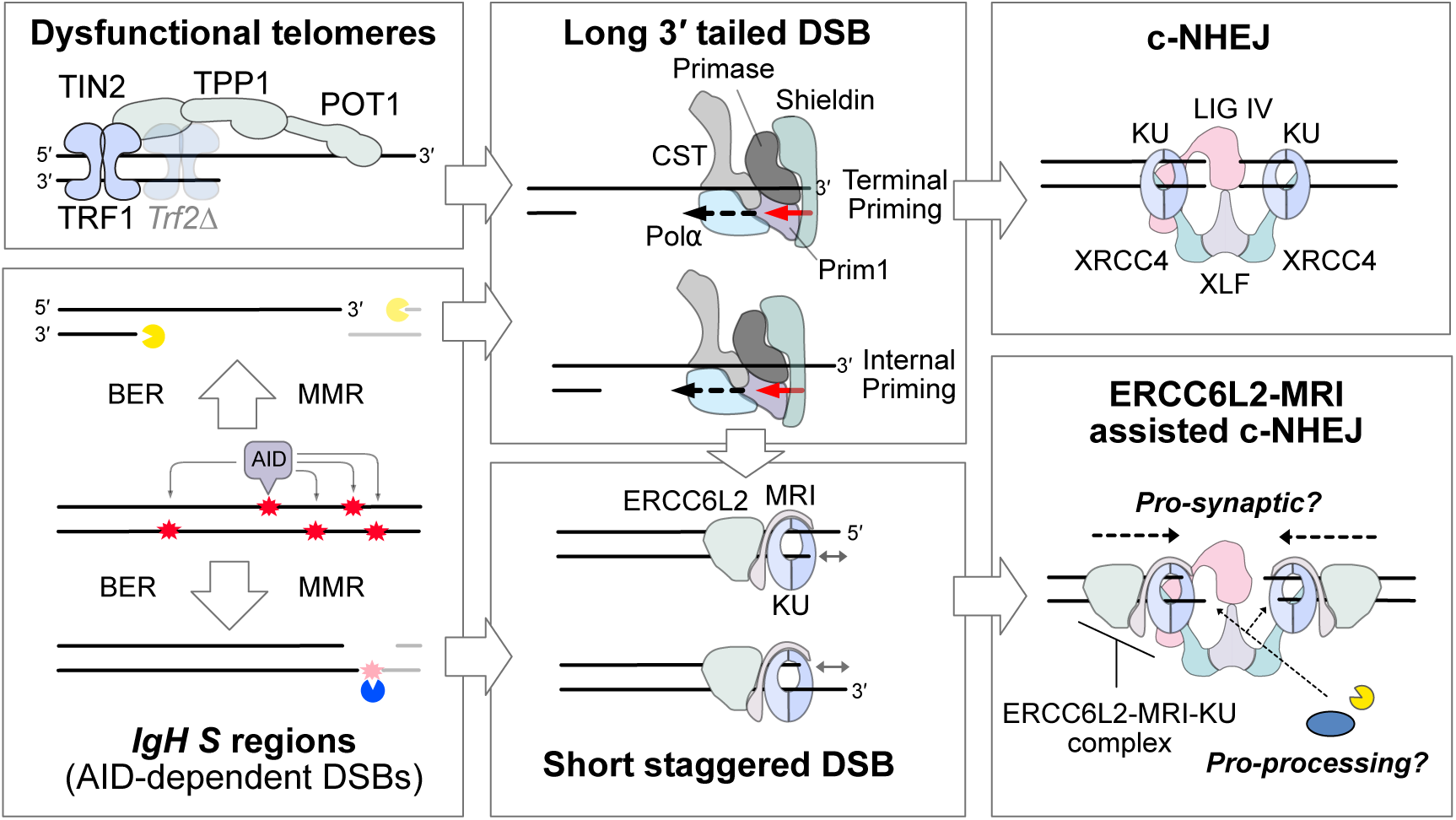
Model of ERCC6L2–MRI–KU–regulated c-NHEJ at staggered DSBs. Depletion of the TRF2 shelterin component exposes DSBs with extended C-strand 3′ overhangs (top left box), while excision and nucleolytic processing of uracil residues (red asterisks) generated by AID-dependent cytosine deamination within *IgH* switch regions result in staggered DSBs of variable overhang length and polarity (bottom left box). In both scenarios, DSBs bearing long 3′ overhangs trigger 53BP1-RIF1-dependent recruitment of shieldin, which co-recruits CST to stimulate 5′-3′ fill-in synthesis, either from DNA termini or from internal positions, thereby generating DSBs with blunt or short-recessed ends. Shieldin-CST-Polα-primase-mediated conversion of blunt DSBs produces ends that are directly ligatable by the short-range c-NHEJ synapsis complex (top right box). By contrast, internally primed fill-in synthesis at telomeric DSBs, or the generation of short-recessed DNA ends by AID, produces intermediates that require ERCC6L2-MRI-dependent coordination of KU at DNA ends (bottom right box) for additional end processing or nucleases, and/or DNA synapsis.

Using a combination of proteomics, structural complex modelling and biochemical model validation, we reveal that ERCC6L2 exists in complex with the KU70/80 heterodimer that is bridged – mostly likely at DNA ends – by MRI, an NHEJ accessory factor that has been attributed contradictory roles in NHEJ stimulation^47^ versus inhibition^44^. Validating the mechanism of ERCC6L2-MRI interaction, which is achieved through an intramolecular β-sheet formed of the HEBO domain in ERCC6L2 and a conserved fold within the core of MRI, we also show that ERCC6L2-MRI co-dependence characterises CSR. Our findings are therefore consistent with a stimulatory physiological role for ERCC6L2-MRI during NHEJ. While ERCC6L2’s precise molecular function is not fully explained by our results, it is tantalising to speculate that it most likely assists in the proper positioning of the KU heterodimer on the staggered DSB through its enzymatic function as a DNA translocase. Such a hypothesized activity may become especially important at complex breaks, such as those formed during CSR at S regions that have a propensity to form stem loops and/or G-quartets due to their GC-rich, repeat-dense nature^50^. If correct, it will remain to be determined if the proper positioning of KU entails pushing the complex towards the ds-ssDNA junction to stimulate short-range fill-in and/or end processing, or towards ssDNA termini, where it might instead favour particular confirmations of the c-NHEJ synaptic complexes that enable internal end-processing events prior to ligation^51,52^ . Indeed, the recent attribution of Polλ-associated pro-insertion activities to ERCC6L2 at Cas9-induced DSBs in the recently published *Repairome*^53^ may add weight to such predictions.

Finally, we speculate that our findings strongly implicate short staggered DSBs of endogenous origins as the likely pathological substrate in ERCC6L2-deficiency. However, considering the scarcity of BMF phenotypes in patients with mutations in other c-NHEJ factors, we speculate it to be unlikely that BMF is precipitated by unrepaired DSBs. Instead, we postulate that ERCC6L2 could perform its stem cell protective function at telomeres. Indeed, we note that there are striking similarities between telomeric ends and AID-dependent DSBs during CSR: both involve dsDNA ends with extended 3’ ssDNA overhangs with propensities to forming secondary structures due to their G-rich content, and ERCC6L2 would not be the only factor to connect the NHEJ machinery to the maintenance of telomeres.

In this light, we highlight our recently demonstration that the DNA-PK complex can be repurposed for DNA end processing and therefore protection at blunt telomeric end intermediates of telomeric leading-strand replication, while its interaction with the shelterin RAP1 counteracts its activity in NHEJ by preventing the action of XRCC4-LIG4^39,42^. Furthermore, whilst the CST complex moonlights at switch regions during CSR, its canonical function is at telomeres, where it bind the ssDNA binding protein POT1 and maintains the C-strand by mediating fill-in synthesis, thereby preventing premature shortening of telomeres^54^. It is therefore tempting to speculate that an analogous end-protective property of the ERCC6L2-MRI-KU - following interactions with telomeric protein that repurpose it to antagonise NHEJ at short-recessed telomeric intermediates of replication and/or repair - could provide a possible explanation for the basis of BMF in ERCC6L2 disease. Such an explanation might even help square with the seemingly opposing role for MRI reported at telomeres, where it was found to inhibit c-NHEJ under certain experimental conditions and/or cell cycle phases^44^. This notion might also help reconcile the inter-species differences in ERCC6L2-dependence in the HSPC niche of mice and men: sufficient age-associated telomere-length attrition might occur during the human lifespan, but is less likely to do so in the C57BL/6-strain derived mice used in this study, whose extensive telomeres limit their suitability as a reliable model of human telomeropathies^55^. Indeed, the fact that replicative lifespan of a stem cell is likely to be limited by the length of the shortest telomere, and not the average telomere length^56,57^, might explain why bulk telomere length measurements in patient samples has thus far failed to detect a role for ERCC6L2 in telomere length homeostasis^2,3^. We suggest using single molecule measurements of telomere length heterogeneity within patient HSCs could represent a fascinating and fertile basis for future investigations into ERCC6L2 disease aetiology.

## Materials & Methods

### Mice

All mice used for this study except for recipient and competitor mice for bone marrow transplantation experiments were generated on a C57BL/6J background. All experiments involved age-matched 8-16-week-old animals on an inbred C57BL/6J background and were approved by the University of Oxford Ethical Review Committee and performed under a UK Home Office Licence in compliance with animal use guidelines and ethical regulations. Animals were house in individually ventilated cages, maintained on a 12h light-dark cycle, and provided with food and water ad libitum.

*53bp1^-/-^* mice (*Trp53bp1^tm1Jc^*, MGI:2654201), *Adh5^-/-^* mice (*Adh5^tm1Stam^*, MGI: 3033711), *Shld2^-/-^* mice (*Shld2^tm1d^*, MGI: 5428631) were generated and described elsewhere^14,24,58^. *Ercc6l2^-/-^* (MGI: 1923501) mice were generated by electroporating Cas9 ribonucleoprotein complexes comprising the following guide RNA (gRNA) spacer sequences (5’-GGTACTTGCGAGATTACCAA-3’ and 5’-ACAGCAGCTTAAAGCGTATA-3’) that target exons 2 and 16, respectively, of the mouse *Ercc6l2* locus into C57BL/6J zygotes. One founder with compound deletions spanning exon 2 to 16 was generated (*Ercc6l2^em1^* 52,455 bp and *Ercc6l2^em2^* 52,601 bp, respectively). Only the *Ercc6l2^em1^*allele was used for this study.

### Competitive bone marrow transplantation

For mixed bone marrow chimera experiments, experiments were performed as described by Wilkinson *et al*.^59^. Briefly, wild-type B6.SJL CD45.1 mice (MGI: 4819849, B6.SJL-Ptprc^a^Pepc^b^/BoyJ) were irradiated with two doses of 5.5 Gy spaced 4 h apart, and subsequently injected with 1 × 10^6^ whole bone marrow cells (approximately 1:1 mixture) from wild-type CD45.1/CD45.2 mice (B6.SJL×C57BL/6) and CD45.2 mice (C57BL/6) with the gene knock-out of interest. Baytril 10% antibiotic solution (BMS, University of Oxford) was added to the drinking water (1.2 mL per 250 mL sterile water) for 4 weeks after irradiation. Donor chimerism was tracked by collecting peripheral blood every 4 weeks for 16 weeks and analysing the ratio of CD45.1/CD45.2 expression via flow cytometry. After 16 weeks, mice were sacrificed and bone marrow donor chimerism was determined by analysing the ratio of CD45.1/CD45.2 expression via flow cytometry. Secondary bone marrow chimera experiments were performed by injecting 1 × 10^6^ pooled whole bone marrow cells from primary recipient into lethally irradiated wild-type B6.SJL C45.1 mice. Mice were treated and donor chimerism was determined as described above.

### Reverse-transcription qPCR

RNA was extracted from mature resting B cells isolated from murine spleens via a RNeasy Mini Kit (Qiagen) with an on-column DNase digestion step as per the manufacturer’s protocol. RNA was then reverse-transcribed using 1 ug of total RNA per sample via the iScript cDNA Synthesis Kit (Bio- Rad). Quantitative PCR reactions were then assembled in 96-well plates using QuantiFast SYBER Green Master Mix (Qiagen) and primer pairs spanning *Ercc6l2* exons 2-3 (5’-AGGGGGCTCAGTTCCTCTAC-3’, 5-GCAAAACTGCAGCCAGAAAT-3), exons 13-14 (5-ACAGCCACAACATGGTTGAA-3, 5’-CGCTGTAGAGCTCACAAACATC-3’) or the *Hprt1*control locus (5’-CTGGTGAAAAGGACCTCTCG-3’, 5’-TGAAGTACTCATTATAGTCAAGGGCA-3’). qPCR was then performed and analysed in a StepOnePlus Instrument (Applied Biosystems) using the “Quantitative - Comparative CT (ΔΔCT)” protocol. Gene expression between samples was normalized using hypoxanthine-guanine phosphoribosyl-transferase (Hprt) as a housekeeping gene. All values were presented as fold-change compared to appropriate controls.

### Flow Cytometry

All flow cytometry experiment were performed using a LSR Fortessa (BD Biosciences), a LSR Fortessa-X20 (BD Biosciences), or Attune NxT flow cytometer (Thermo Fisher). Analysis was performed using FlowJo (version 10.10.0, BD Bioscience). FACS buffer used in all experiments was 2% bovine serum albumin (Sigma) in PBS with 0.025% NaN_3_ (Sigma).

### Haematopoietic stem and progenitor cells

Briefly, mice were sacrificed and one femur and tibia were dissected out. Bone marrow was collected by flushing the bones with PBS using a 23-gauge needle. The cell suspension was then counted using a Pentra ES60 Hematology Analyzer (Horiba) and 5−10 × 10^6^ cells were stained with 300 μL of the biotin-conjugated lineage antibody stain (see Table 1) for 30 min at 4°C, washed once with PBS, and then stained with 300 μL of the secondary antibody stain (see Table 2) for 90 min at 4°C, and washed once in PBS. Subsequently, cells were resuspended in FACS buffer with propidium iodide (0.5 μg/mL, Sigma) and analysed on the flow cytometer.

**Table 2:** List of antibodies for haematopoietic stem and progenitor analysis. Working dilutions are listed.

| Antibody | Manufacturer | Catalogue number | Dilution |
| --- | --- | --- | --- |
| c-Kit-APC | Invitrogen | 17-1171-83 | 1:300 |
| CD150-PE/Cy7 | BioLegend | 115914 | 1:300 |
| CD34-FITC | invitrogen | 11-0341-85 | 1:75 |
| CD48-BV421 | BioLegend | 103427 | 1:300 |
| Sca1-PE | BioLegend | 122508 | 1:300 |
| Streptavidin-APC-eFluor780 | Invitrogen | 47-4317-82 | 1:300 |

### Peripheral blood chimerism

Flow cytometry was performed as described by Wilkinson *et al.*^59^ . Briefly, 50 μL of peripheral blood was collected from the tail vein of recipient mice into Lithium heparin-coated tubes (Sarstedt). Erythrocytes were subsequently lysed for 15 min at room temperature using ACK lysis buffer (ChemCruz). Lysis was repeated once, then the remaining mononuclear cells were incubated in 100 μL of antibody stain (see Table 3) for 30 min at 4°C. Cells were twice washed in FACS buffer, resuspended in FACS buffer with propidium iodide (0.5 μg/mL, Sigma) and analysed on the flow cytometer.

**Table 3:** List of antibodies for analysis of peripheral blood chimerism. Working dilutions are listed.

| Antibody | Manufacturer | Catalogue number | Dilution |
| --- | --- | --- | --- |
| B220-APC-eFluor780 | invitrogen | 47-0452-82 | 1:1000 |
| CD11b-PE | invitrogen | 12-0112-82 | 1:2000 |
| CD4-APC | invitrogen | 17-0042-83 | 1:2000 |
| CD45.1-PE/Cy7 | BioLegend | 110730 | 1:400 |
| CD45.2- eFluor450 | invitrogen | 48-0454-82 | 1:333 |
| CD8-APC | invitrogen | 17-0081-83 | 1:2000 |
| Ly6G-PE | invitrogen | 12-5931-82 | 1:2000 |

### Micronucleus assay

Micronucleus levels in peripheral blood were analysed as described by Balmus *et al.*^17^. Briefly, Briefly, 50 μL of peripheral blood were collected from the tail vein into a heparin solution and immediately fixed using cold methanol. Samples were stored at -80°C until further processing. Cells were stained with CD71-FITC (1:400, BioLegend, cat. no. 113806) for 60 min at 4°C and ultimately resuspended in FACS buffer with propidium iodide (0.5 μL, Sigma).

### B cell analysis and flow cytometry

Mice were sacrificed, and the spleen, and one femur and tibia were dissected out. Bone marrow was collected by flushing the bones with PBS using a 23-gauge needle. The spleen mashed into a single-cell suspension. The cell suspensions were then counted using a Pentra ES60 Hematology Analyzer (Horiba) and stained with 50 μL of the antibody stain (see Tables 4 and 5) for 20 min at 4°C. Cells were washed twice in FACS buffer, then resuspended in FACS buffer and analysed on the flow cytometer.

**Table 4:** List of antibodies for analysis of B cell development analysis in the bone marrow. Working dilutions are listed.

| Antibody | Manufacturer | Catalogue number | Dilution |
| --- | --- | --- | --- |
| ZombieAqua | BioLegend | 423102 | 1:200 |
| B220-BV605 | BioLegend | 103244 | 1:200 |
| IgM-FITC | BioLegend | 406506 | 1:200 |
| BP-1-PE | BioLegend | 108308 | 1:200 |
| CD43- PerCP/Cy5.5 | BD Biosciences | 562865 | 1:200 |
| CD24-AlexaFluor700 | BioLegend | 101836 | 1:500 |
| IgD-APC/Cy7 | BioLegend | 405716 | 1:500 |
| FC-block | BD Biosciences | 553141 | 1:500 |

**Table 5:** List of antibodies for analysis of B cell development analysis in the spleen. Working dilutions are listed.

| Antibody | Manufacturer | Catalogue number | Dilution |
| --- | --- | --- | --- |
| ZombieAqua | BioLegend | 423102 | 1:200 |
| B220-BV605 | BioLegend | 103244 | 1:200 |
| IgM-PacificBlue | invitrogen | 48-5890-82 | 1:200 |
| IgD-FITC | BioLegend | 406506 | 1:200 |
| CD23-PE | BD Biosciences | 553139 | 1:200 |
| CD19-PerCP/Cy5.5 | BioLegend | 115534 | 1:500 |
| CD93-APC | BioLegend | 136510 | 1:100 |
| CD21-APC/Cy7 | BioLegend | 123418 | 1:200 |
| FC-block | BD Biosciences | 123418 | 1:500 |

### T cell analysis and flow cytometry

Mice were sacrificed and thymus was dissected out and mashed into a single-cell suspension. The cell suspension was then counted using a Pentra ES60 Hematology Analyzer (Horiba) and stained with 50 μL of the antibody stain listed in Table 6 for 20 min at 4°C. Cells were washed twice in FACS buffer, then resuspended in FACS buffer and analysed on the flow cytometer.

**Table 6:** List of antibodies for analysis of T cell development. Working dilutions are listed.

| Antibody | Manufacturer | Catalogue number | Dilution |
| --- | --- | --- | --- |
| ZombieAqua | BioLegend | 423102 | 1:200 |
| TCR-β-BV605 | BioLegend | 109241 | 1:200 |
| CD44-FITC | BioLegend | 103022 | 1:200 |
| CD25-PE | BioLegend | 102008 | 1:500 |
| CD8a-PerCP | BioLegend | 100732 | 1:500 |
| CD4-AlexaFluor700 | BioLegend | 100430 | 1:500 |
| CD3-APC/Cy7 | BioLegend | 100248 | 1:100 |
| FC-block | BD Biosciences | 553141 | 1:500 |

### *Ex vivo* splenic B cell culture, stimulation and flow cytometry

Murine splenic B cells were isolated from erythrocyte-lysed single-cell suspensions by magnetic negative selection using a B cell isolation kit (Miltenyi Biotec, 130-090-862) as per manufacturer’s instructions. Isolated B cells were then seeded at 3 × 105 cells/mL in RPMI-1640 medium (Sigma) supplemented with 10% FBS (Gibco), 1%Penicillin-Streptomycin (10,000U/mL, Sigma), 2mM L-Glutamine (Gibco), 1X MEM nonessential amino acids (Gibco), 1mM sodium pyruvate (Gibco), and 50μM β-mercaptoethanol at 37°C and 5% CO2 in a humidified atmosphere. B cells were stimulated with LPS (Sigma, 5ug/mL), mouse recombinant IL-4 (10ng/mL, PeproTech), and agonist anti-CD40 antibody (0.5ug/mL, Miltenyi Biotec). Four days after seeding, cells were collected and stained with 50 μL of the primary antibody stain (see Table 7) for 20 min at 4°C, washed twice in FACS buffer, and then stained with 50 μL of the secondary antibody stain (see Table 8) for 20 min at 4°C. Cells were washed twice with FACS buffer, and on the flow cytometer. Cell proliferation was assessed using Cell Trace Violet (invitrogen) according to manufacturer’s instructions.

**Table 7:** List of primary antibodies for B cell isotype switching analysis. Working dilutions are listed.

| **Antibody** | **Manufacturer** | **Catalogue number** | **Dilution** |
| --- | --- | --- | --- |
| IgG1-biotin | BD Biosciences | 553441 | 1:100 |
| IgG2b-biotin | BD Biosciences | 406704 | 1:100 |
| IgG3-biotin | BD Biosciences | 553401 | 1:100 |
| FC-block | BD Biosciences | 553141 | 1:500 |

**Table 8:** List of secondary antibodies for B cell isotype switching analysis. Working dilutions are listed.

| **Antibody** | **Manufacturer** | **Catalogue number** | **Dilution** |
| --- | --- | --- | --- |
| ZombieAqua | BioLegend | 423102 | 1:200 |
| Streptavidin-APC | Invitrogen | 17-4317-82 | 1:500 |
| IgE-PE | BioLegend | 406908 | 1:200 |

**Table 9:** List of primary and secondary antibodies used for Western blotting. Working dilutions are listed.

| Antibody | Manufacturer | Catalogue number | Dilution |
| --- | --- | --- | --- |
| α-TUBULIN | Sigma-Aldrich | 00020911 | 1:10,000 |
| CHK2 | Sigma-Aldrich | 05-649 | 1:1000 |
| FLAG-M2 | Sigma-Aldrich | F1804 | 1:2000 |
| HA-tag | Cell Signalling | 3724 | 1:1000 |
| KU70 | Cell Signalling | 4588 | 1:1000 |
| anti-mouse IgG-HRP | Invitrogen | 31432 | 1:50,000 |
| anti-rabbit IgG-HRP | Invitrogen | 31462 | 1:50,000 |
| Mouse TrueBlot ULTRA | Rockland | 18-8817-30 | 1:5000 |

### CSR assay in CH12-F3 cells

CH12-F3 cells were plated at 1.5x10^5^ cells/mL in complete RPMI-1640 medium supplemented with anti-CD40 antibody (1μg/mL, Miltenyi Biotec), IL-4 (20ng/mL, PeproTech), and TGF-β (10ng/mL, PeproTech) to stimulate isotype switching from IgM to IgA. 48h after stimulation, cells were assayed for class switching by flow cytometry using IgA-PE (1040-09, SouthernBiotech, 1:250) antibody.

### V(D)J reporter assay in *v-abl* cells

*v-abl* pre-B cells were seeded at a final concentration of 1x10^6^ cells/mL in complete v-Abl media and treated with 3 µM imatinib (13139-25, Cambridge Bioscience) for 96h. Cells were then collected, washed once in FACS buffer, and stained with 50 µL of antibody stain containing hCD4-AF647 (1:50, 300520, BioLegend) and ZombieAqua (1:250, 423102, BioLegend) for 20 min at 4° C. Cells were twice washed with FACS buffer, then resuspended in FACS buffer and analysed on the flow cytometer.

### Cell lines and culture conditions

Human embryonic kidney 293T cells (HEK293T) and mouse embryonic fibroblasts (MEF) were cultured in DMEM (Sigma) supplemented with 10% FBS (Gibco), 1% Penicillin-Streptomycin (10,000U/mL, Sigma), and 2 mM L-Glutamine (Gibco) at 37°C and 5% CO_2_ in a humidified atmosphere. E12.5 MEFs were prepared and SV40 large T antigen-immortalized as per standard procedure.

CH12-F3 cells were cultured in RPMI-1640 medium (Sigma) supplemented with 10% FBS (Gibco), 5% NCTC-109 medium (Sigma), 1% Penicillin-Streptomycin (10,000 U/mL, Sigma), 2 mM L-Glutamine (Gibco), and 50 μM β-mercaptoethanol (Gibco) at 37°C and 5% CO2 in a humidified atmosphere.

Viral Abelson kinase (v-abl) transformed pre-B cell lines were cultured in RPMI-1640 medium (Sigma) supplemented with 10% FBS (Gibco), 1% Penicillin-Streptomycin (10,000 U/mL, Sigma), 2mM L-Glutamine (Gibco), and 100 μM β-mercaptoethanol (Gibco) at 37°C and 5% CO_2_ in a humidified atmosphere.

### CRISPR/Cas9 genome editing

*Ercc6l2^-/-^* CH12-F3 cells were generated using CRISPR/Cas9 as previously described^48^. For all cell lines generated for this study, gene-specific sgRNAs comprising the following spacwer sequences were cloned into pX458 vectors (Addgene #48138): sgMri 5’-agaggggaggaccctcgttt-3’ (gene knockout); sgXrcc4 5’-tttgttattacacttactga-3’ (gene knockout); sgErcc6l2 5’-ggagtctgcgcgggactgcg-3’ (gene editing – 5’-tagging). Cells were then electroporated with the plasmid using the Lonza 4D-Nucelofector according to the manufacturer’s protocol for each cell line. 24h post nucleofection, GFP-positive cells were sorted using a Sony SH800 cell sorter, with the brightest 10% being pooled for subsequent recovery. To achieve isogenic cell lines, the cells were then seeded at 0.5 cells/well in 96-well plates and individual clones were isolated and screened after 7 days of outgrowth. Clones bearing bi-allelic indel mutations were identified using native PAGE resolution of PCR amplicons of the edited loci and gene disruption was subsequently confirmed via Sanger sequencing.

### Lentivirus generation and infection

To generate complemented cell lines expressing MRI, point mutants of MRI, or the corresponding transduction control (GFP), lentiviral transductions were performed. HEK293T cells were co-transfected with second-generation packaging plasmids and pLenti-PGK-PURO-DEST vectors (Addgene #19068) containing cloned transgene inserts. Viral supernatants were collected 48 h and 72h post transfection. Typically, cells were then spinoculated with viral supernatants supplemented with polybrene (4μg/mL, Merck) and HEPES (20mM, Gibco) at 1500rpm for 90 min at 30°C. Stable cell lines were subsequently selected and maintained by addition of puromycin (500ng/mL, Gibco).

### Cell survival assay

CH12-F3 or MEF cells were seeded in triplicate per drug concentration in 96-well plates at a density of 1250 or 600 cells per well, respectively. Then, etoposide or cisplatin (Merck) was added to the final indicated concentrations. Four (CH12-F3) or seven (MEF) days after cell seeding, growth medium with resazurin was added to a final concentration of 10 ug/mL resazurin. Plates were then returned to the incubator for 2-4h or until the growth medium in the untreated control wells developed a pink colour. Relative fluorescence was measured with a BMG LABTECH CLARIOStar plate reader. After subtraction of the background fluorescence, relative survival compared to the untreated control was calculated.

### Proteomics and mass spectrometry

Cell pellets were collected from cultures of 1.4 × 108 CH12-F3 cells ±10 Gy irradiation and 60 min recovery. Cells were lysed in BLB (benzonase lysis buffer: 100 mM NEM, 40 mM KCl, 2 mM MgCl2, 10% glycerol, 1% NP-40, 50 U/mL benzonase (Millipore), 1X cOmplete protease inhibitor cocktail (Roche), 0.5% (v/v) protease inhibitor cocktail 2 and 3 (Sigma-Aldrich)) and incubated for 10 min on ice. The salt concentration was then adjusted to 450 mM KCl and lysates were further incubated on wheel for 30 min at 4°C. Subsequently, extracts were clarified by centrifugation at 17,000 G for 15 min and then diluted in NSB (no-salt buffer: 100 mM NEM, 20mM HEPES, pH 7.9, 0.5 mM DTT, 0.5 mM EDTA, 10% glycerol, 1X cOmplete protease inhibitor cocktail (Roche), and 0.5% (v/v) protease inhibitor cocktail 2 and 3 (Sigma-Aldrich)) to reach a final salt concentration of 125 mM KCl. The clarified lysate was incubated with MagStrep-XT magnetic beads (iba) for 4 h at 4°C, subsequently washed in wash buffer (100 mM KCl, 20 mM HEPES, pH 7.9, 0.5 mM DTT, 10% glycerol, 0.05% NP-40, 1X cOmplete protease inhibitor cocktail (Roche), and finally eluted in 1X BXT buffer (iba) supplemented with 1X cOmplete protease inhibitor cocktail (Roche). An aliquot was mixed with Laemmli buffer and boiled at 95°C for 5 min for validation using Western blot. The remaining eluates were snap-frozen for further downstream processing for proteomic analysis.

The eluted complexes were solubilized in 4% (w/v) SDS and 50 mM TEAB pH8.5, reduced and alkylated using DTT and iodoacetamide and digested with trypsin on S-Trap microcolumns as per the manufacturer’s instructions. Peptides were subjected to LCMS/MS using an Ultimate 3000 RSLC coupled to an Orbitrap Fusion Lumos Tribrid mass spectrometer with Easyspray source and FAIMS interface, (Thermo Fisher Scientific). Peptides were injected onto a reverse-phase trap (Pepmap100 C18 100μm x 2cm) for pre-concentration with loading buffer (100% water, 0.05% TFA), at 5 μL/min for 10 minutes. The peptide trap was then switched in-line with the analytical column (Easy-spray Pepmap RSLC C18 2um, 15cm x75um ID). Peptides were eluted from the column using a linear solvent gradient at a flow rate of 300nl/min with the following gradient: linear 2–35% of buffer B (Mobile phase B: 80% acetonitrile, 20% water, 0.1% formic acid, Mobile phase A: 100% water, 0.1% formic acid) over 44 min, sharp increase to 95% buffer B within 0.5min, isocratic 95% of buffer B for 4.5 min, sharp decrease to 2% buffer B within 1.1min and isocratic 2% buffer B for 5 min. The FAIMS interface was set to -45V and -65V at standard resolution. The mass spectrometer was operated in DDA positive ion mode with a cycle time of 1.5sec. The Orbitrap was selected as the MS1 detector at a resolution of 60000 with a scan range of m/z 375 to 1500. RF-lens was set to 30%. Peptides with charge states 2 to 5 at a minimum intensity threshold of 5.0e3 were selected for fragmentation in the ion trap using HCD as collision energy. Once selected, a dynamic exclusion for 45sec with a 10ppm mass tolerance was applied. A dependent scan was only performed on the single charge state per precursor. The Ion Trap was selected for data dependent MS2 fragmentation with an isolation window of m/z 1.2 and no isolation offset. The peptides were fragmented with fixed HCD as activation type at 28% HCD Collision Energy. The Ion Trap scan rate was set to Rapid with a maximum injection time of 45ms. The raw data files were converted into mgf using MSconvert (ProteoWizard) and searched using Mascot (version 2.8.1, Matrix Science) with trypsin as the cleavage enzyme, carbamidomethylation on cysteines as fixed modification and oxidation of methionines as a variable modification against the Uniprot mouse database (downloaded on 28Oct2024, containing 54747 protein sequences). The mass accuracy for the MS scan was set to 20ppm and for the fragment ion mass to 0.6Da.

Scaffold (version 5.3.3, Proteome Sofwarte Inc.) was used to validate MS/MS based peptide and protein identification. Peptide identifications were accepted if they could be established at >95.0% probability by the Peptide Prophet algorithm^60^ with Scaffold delta-mass correction. Protein identifications were accepted if they could be established at >99.0% probability and contained at least 1 identified peptide. Protein probabilities were assigned by the Protein Prophet algorithm^61^.

### AlphaFold modelling

The structures of ERCC6L2- MRI-Ku70-Ku80 were modelled using Alphafold 3^43^. The sequences of *homo sapiens* ERCC6L2, MRI, Ku70, Ku80, along with DNA substrates consisting of two 30nt complementary strands with various length of single strand overhangs, were used as input into the AlphaFold 3 web server (https://alphafoldserver.com/) for protein complex prediction. The resulting structural models were assessed based on the per-residue confidence metric(plDDT), and the model with the highest plDDT score was selected for further analysis. Subsequent structural analysis is carried out at regions with high confidence (plDDT>70). As for domain interface evaluation, only high confidence regions with accurate relative domain position (Expected Position Error <15) were considered. To ensure the prediction and interpretation was reliable and robust, additional structural alignments and comparisons were preformed among all AF3 models; MRI and Ku80 interface was superimposed with the reported crystal structure (PDB: 6TYU).

### SDS-PAGE-Western Blot

Protein lysates were mixed with Laemmli buffer and boiled at 95°C for 5 min before loading on NuPAGE 4-12% Bis-Tris 1.0 mm polyacrylamide gels (invitrogen). SDS-PAGE gels were run for at 95V for 15min, followed by 150V for 60min before transferring to 0.45-μm nitrocellulose membranes (GE Healthcare) at 50 mA overnight. After transfer, membranes were blocked for 30 min with 5% milk in PBS-T and then incubated with primary antibodies listed in Table 1.10 in 5% BSA in PBS-T with 0.025% NaN_3_ overnight. On the next day, membranes were incubated with secondary antibodies conjugated to horseradish peroxidase (HRP) for at least 1 h at room temperature. All antibodies are listed in Table 8. Membranes were developed using Clarity Western enhanced chemiluminescence substrate (Bio-Rad) or SuperSignal West Pico Plus (Thermo Fisher) and imaged using a ChemiDoc MP (Bio-Rad). Images were analysed using Image Lab (Bio-Rad).

### Plasmids

For the dysfunctional telomeres c-NHEJ assay, sgRNAs against *Apollo* (target: 5’-GTGATGGGAGAGCAGTAGAG-3’) in lentiCRISPR v2 hygro (Addgene plasmid #98291, a gift from B. Stringer^62^ and against *Rap1* (target: 5′-GCAGTCTAGGATGTACTGCG-3) in lentiCRISPR v2 puro (Addgene plasmid #52961, a gift from F. Zhang^63^ have been previously described^39,42^. The sgRNA against mouse *Trf2* (target: 5’- CAGATCCGGGACATCATGCA-3’) was cloned in lentiCRISPR v2 Puro.

### Transfection and transduction

Lentiviral particles were obtained after transfection of 293 T cells (ATCC, CRL-3216), using CaPO_4_ precipitation with 10 µg of lentiviral vector plasmid and a total of 10 µg of packaging plasmids. The viral supernatant was filtered through a 0.45 μm filter, supplemented with 4 μg/ml polybrene, and used to transduce the target cells with three infections per day (6–12 h intervals) over 2 days. For *Trf2* deletion, transduced MEFs were selected for 2 days in 2–4 µM puromycin (t=0 at 12 hours after the first infection). For *Rap1* and *Apollo* deletions, MEFs transduced first with sgRap1 were selected for 2 days in 340 µM hygromycin before transduction with sgApollo and selection for 2 days in 2–4 µM puromycin (t=0 at 12 hours after the first infection with sgApollo).

### Immunoblot

Cells were harvested via trypsinization, washed in PBS and lysed in 2× Laemmli buffer. After denaturation for 10 min at 95°C and chromatin shearing the chromatin with sonication at 40% amplitude for 15 s, 5 s ON and 5 s OFF (Fisherbrand sonicator, Model; FB705; power 700 W; 2000 Park Lane, Pittsburgh, PA, 15275). The lysate was resolved via SDS/PAGE and transferred to a nitrocellulose membrane. Western blot was performed with 5% milk in PBS containing 0.1% (v/v) Tween-20 (PBS-T) using the following antibodies: β-actin (#3700; Cell Signaling), RAP1 (#5433, Cell Signalling) and TRF2 (#13136, Cell Signaling), followed by goat anti-rabbit (31460, Invitrogen) or anti-mouse (31430, Invitrogen) IgG–HRP peroxidase secondary antibody. Signals were detected according to the manufacturer’s instructions using chemiluminescence western blotting detection reagents (Cytiva) on ChemiDoc (Bio-Rad) imaging systems.

### Fluorescence in situ hybridization (FISH)

For telomere FISH, MEFs treated with 0.2 μg/ml Colcemid in the last 1–2 h before collection by trypsinization. Harvested cells were swollen in a hypotonic solution of 0.055–0.075 M KCl at 37°C for 15–30 min before fixation in methanol/acetic acid (3:1) overnight at 4°C. Cells were dropped onto glass slides and allowed to age overnight before dehydration through an ethanol series of 70%, 95% and 100%. Telomere ends were hybridized with Alexa Fluor 488-OO-(TTAGGG)3 or Alexa Fluor 647-OO-(CCCTAA)3 in hybridization solution (70% formamide, 1 mg/ml blocking reagent (1109617601, Roche), and 10 mM Tris-HCl pH 7.2) for 2 h followed by two washes in 70% formamide; 0.1% Bovine Serum Albumin (BSA); 10 mM Tris-HCl, pH 7.2 for 15 min each, and two washes in 0.08% Tween-20, 0.15 M NaCl, 0.1 M Tris-HCl, pH 7.2 or PBS for 5 min each. Chromosomal DNA was counterstained with the addition of 4′,6-diamidino-2-phenylindole (DAPI) (D1306, Invitrogen) to the second wash. Slides were left to air-dry and mounted in antifade reagent (Prolong Gold Antifade P36934, Fisher).

Images were acquired on a DeltaVision RT microscope system (Applied Precision) with a PlanApo 60 × 1.40 NA objective lens (Olympus America, Inc.) at 1 × 1 binning and multiple 0.2 μm Z-stacks using SoftWoRx software and deconvolved. 2D-maximum intensity projection images were obtained using SoftWoRx software. Chromatid and chromosome-type fusions were analysed using Fiji software^64^ .

### Statistics

Prism 10 (version 10.5.0) was used for graph generation and statistical analysis. The relevant statistical methods for individual experiments are listed in the figure legends. For FISH analysis, at least ten metaphases per condition were scored for each experiment. Significance was assessed by calculating the p-value using one-way ANOVA Turkey’s multiple comparisons test. P-values ≤ 0.05 were considered statistically significant.

## Acknowledgments

We thank all members of the laboratory of J.R.C. and S. Rottenberg (University of Berne) for discussions. K.J. Patel (University of Oxford) for *Adh5* mutant mice, and S. Zha (Columbia University), B. Sleckman and B.-R. Chen (University of Alabama at Birmingham) for advice, reagents, protocols and cell lines. The University of Oxford Department of Biomedical Services (BMS) Functional Genetics and JR Hospital BMS facilities for technical support. This work was funded by the Medical Research Council (MRC, UK) project grant MR/R017549/1 and MRC Molecular Haematology Unit grants MC_UU_00016/19 and MC_UU_00029/2. Salary to J.R.C. and funding for his group comes from a Cancer Research UK (CRUK) Senior Cancer Research Fellowship (RCCSCF-Nov21\100004) which currently provides salary support to P.I.R. and J.S.M.. A.K. was previously supported by EMBO Long-Term (EMBO ALT 542-2020). P.I.R.’s DPhil in Medical Sciences studentship (2021-2025) was co-supported through funding from Radcliffe Department of Medicine (University of Oxford) and the Lister Institute of Preventative Medicine. Y.S is supported by the Breast Cancer Now Project Grant (2022.11PR1585). J.R.C. was the former recipient of Lister Institute Research Prize Fellowship funding. C.S. is partially supported by the Lions forskningsfond. F.L. is supported by Cancerfonden (23 3038 Pj) and Vetenskapsrådet (2021-02788) grants and the Knut and Alice Wallenberg Foundation. A.C.W. was funded by the Kay Kendall Leukaemia Fund and the Wellcome Trust.

## Author Contributions

Conceptualization, J.R.C.. Methodology, P.I.R., C.S., Y.S., A.K., J.S.M., B.D., A.C.W., F. L. & J.R.C.. Investigation, P.I.R., C.S., Y.S., A.K., J.S.M., B. D. & J.R.C.. Writing, original draft, P.I.R. & J.R.C.. Writing, editing, P.I.R., Y.S., F.L. & J.R.C.. Funding Acquisition, A.C.W., F.L & J.R.C. Supervision, A.C.W., F.L & J.R.C.

## Competing Interest Statement

The authors declare no competing interests.

Correspondence and requests for materials should be addressed to J. Ross Chapman.

