## Extended Data Figures for "The ERCC6L2-MRI-KU complex coordinates NHEJ at staggered DNA double-strand breaks"

**Affiliations:**
1. Genome Integrity laboratory, Medical Research Council Molecular Haematology Unit, MRC Weatherall Institute of Molecular Medicine, Radcliffe Department of Medicine, The University of Oxford, Oxford, UK.
2. Department of Biomedical and Clinical Sciences, Division of Molecular Medicine and Virology, Faculty of Medicine and Health Sciences, Linköping University, Linköping, Sweden.
3. Francis Crick Institute, 1 Midland Rd, London, UK.
4. Department of Haematology, Cambridge Stem Cell Institute, University of Cambridge, Cambridge, UK

**Corresponding author:**
*


The PDF file contains:

Extended Data Figures 1-9

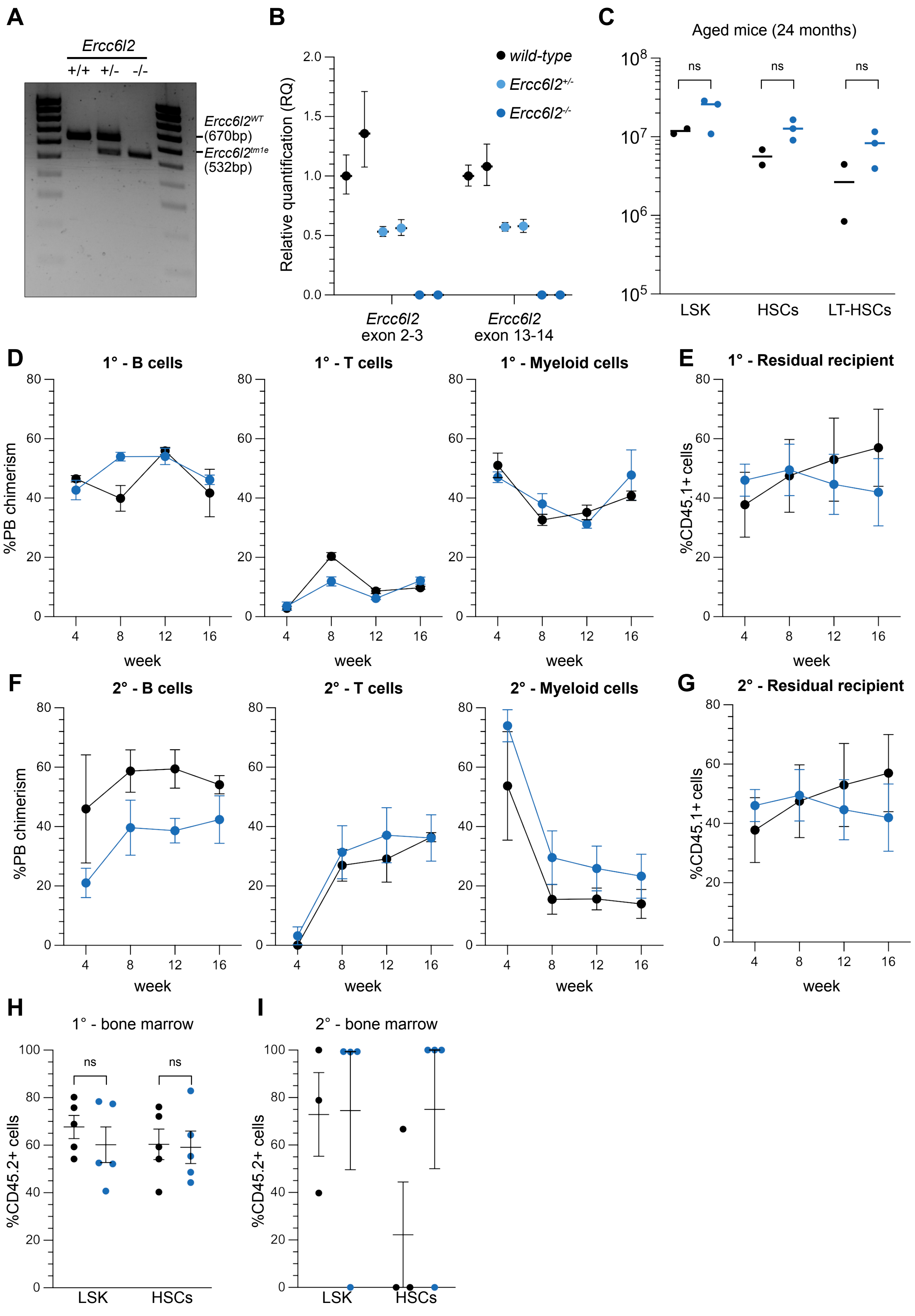


### Extended Data Fig 1

**Extended Data Fig. 1. Resilient Haematopoiesis characterises a mouse model of ERCC6L2 disease**

**A.** Representative image of PCR amplicons from genomic DNA from ear biopsies from mice of the indicated genotype. Bands of different size correspond to *Ercc6l2*$^{em1}$ allele (532bp), or *WT* allele (670bp).

**B.** RT-qPCR result of *Ercc6l2* transcript expression in splenic B cells isolated from mice of the indicated genotypes. Gene expression was normalized to *Hprt1* (*n*=2 per genotype, where each data point represents one mouse).

**C.** Absolute number of Lin$^-$ Sca1$^+$ c-Kit$^+$ (LSK) cells, CD34$^-$ LSK cells (HSCs) and CD34$^-$ CD48$^-$ CD150$^+$ LSKs (-HSCs) in the bone marrow of mice aged for 24 months (one femur and one tibia) (*n*= 2-3 mice per genotype, where each data point is a single mouse). Significance was determined by an unpaired two-tailed *t*-test (mean ± SEM). ns, not significant

**D.** Plots show donor vs. competitor chimerism in mononucleated peripheral blood (PB) during the primary bone marrow transplant. B cells are identified as B220$^+$, T cells as CD4$^+$ and/or CD8$^+$, and myeloid cells as CD11b$^+$ and/or Ly6G$^+$ (*n*= 4-5 mice per test condition, mean ± SEM).

**E.** Plot shows residual recipient chimerism (CD45.1$^+$) in mononucleated peripheral blood during the primary bone marrow transplant (*n*= 4-5 mice per test condition, mean ± SEM).

**F.** Plots show donor vs. competitor chimerism in mononucleated peripheral blood (PB) during the secondary bone marrow transplant. B cells are identified as B220$^+$, T cells as CD4$^+$ and/or CD8$^+$, and myeloid cells as CD11b$^+$ and/or Ly6G$^+$ (*n*= 3-4 mice per test condition, mean ± SEM).

**G.** Plot shows residual recipient chimerism (CD45.1$^+$) in mononucleated peripheral blood during the secondary bone marrow transplant (*n*= 4-5 mice per test condition, mean ± SEM).

**H.** Plots show donor vs. competitor chimerism in the bone marrow in the indicated cell types during the primary bone marrow transplant (*n*= 5 mice per test condition, mean ± SEM).

**I.** Plots show donor vs. competitor chimerism in the bone marrow in the indicated cell types during the secondary bone marrow transplant (*n*= 3-4 mice per test condition, mean ± SEM)

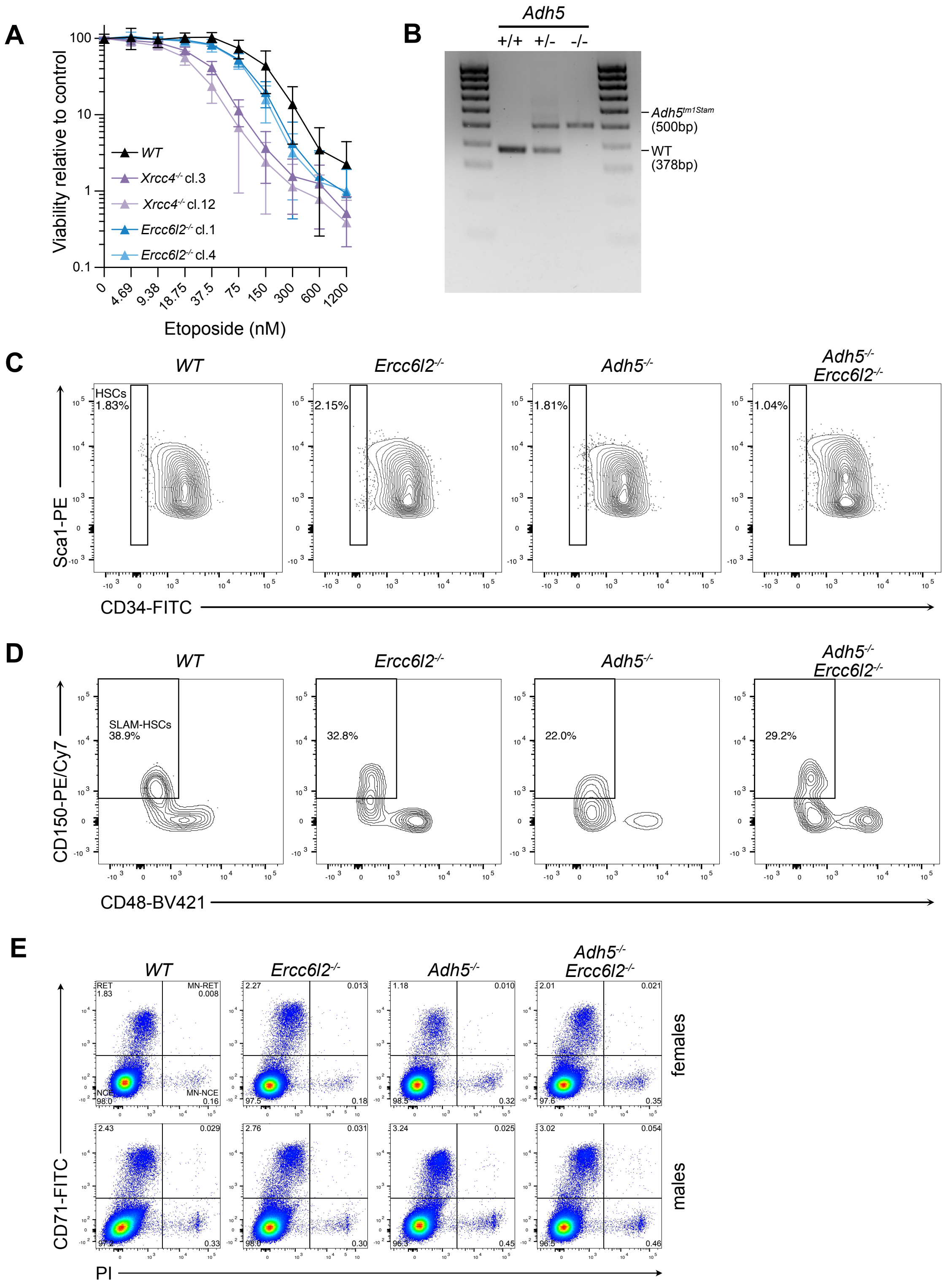


### Extended Data Fig 2

**Extended Data Fig. 2. Combined loss of *Ercc6l2* and *Adh5* does not lead to synergistic loss of haematopoietic stem cells**

**A.** Survival of the indicated MEF cell lines grown for 7 days in the presence of indicated doses of etoposide (*n*= 3 biological replicates, with 3 technical replicates each, mean ± SD).

**B.** Representative image of PCR amplicons from genomic DNA from ear biopsies from mice of the indicated genotype. Bands of different size correspond to $Adh5^{tm1Stam}$ allele (500 bp), or *WT* allele (378bp).

**C.** Representative flow cytometry plots showing LSK cells used to quantify HSC frequencies in the bone marrow.

**D.** Representative flow cytometry plots showing $CD34^-$ LSK cells used to quantify SLAM-HSC frequencies in the bone marrow.

**E.** Representative flow cytometry plots showing peripheral blood used to quantify micronucleated erythrocytes. RET, reticulocytes; MN-RET, micronucleated reticulocytes; MN-NCE, micronucleated normochromatic erythrocytes; NCE, normochromatic erythrocytes

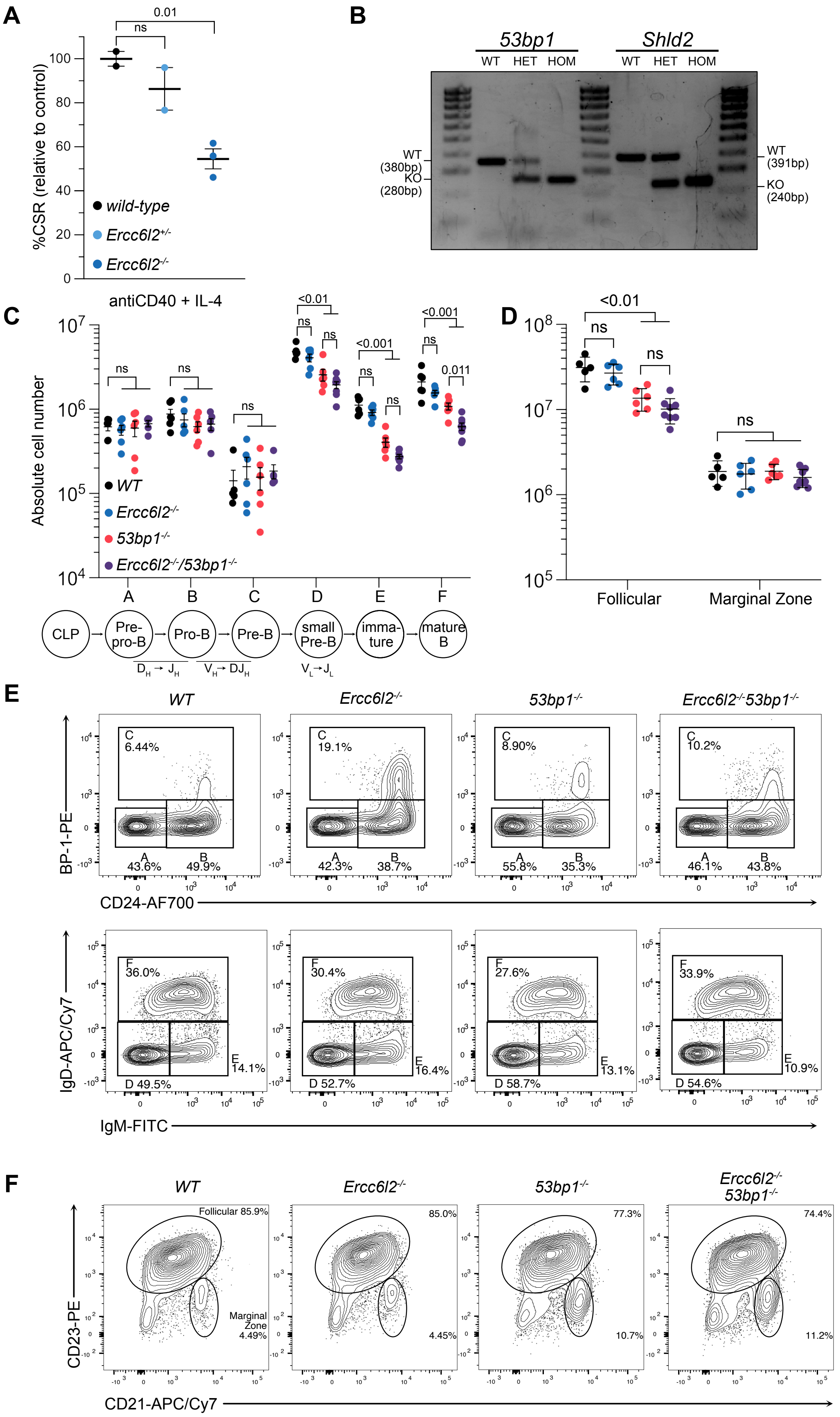


### Extended Data Fig 3

**Extended Data Fig. 3. B cell development in *53bp1$^{-/-}$ Ercc6l2$^{-/-}$* mice**

**A.** Splenic B cells cultured with indicated cytokines for 96 h and stained for IgG1 (*n*=2-3 mice per genotype, where each data point represents a single mouse). Isotype switching frequency was normalized to two wild-type mice.

**B.** Representative image of PCR amplicons from genomic DNA from ear biopsies from mice of the indicated genotype. Bands of different size correspond to *Trp53bp1$^{tm1Jc}$* allele (280bp), or *WT* allele (380bp), and *Shld2$^{tm1d}$* allele (240bp) or *WT* allele (391bp).

**C.** Absolute number of B220$^{+}$ B cell precursors in the bone marrow (one femur and one tibia) *n*= 5-8 mice per genotype, where each data point is a single mouse). Significance was determined by an unpaired two-tailed *t*-test (mean ± SEM). ns, not significant; A, B220$^{+}$CD43$^{+}$BP1$^{-}$CD24$^{-}$; B, B220$^{+}$ CD43$^{+}$ BP1$^{-}$CD24$^{+}$; C, B220$^{+}$ CD43$^{+}$ BP1$^{+}$ CD24$^{+}$; D, B220$^{+}$ CD43$^{-}$ IgM$^{-}$ IgD$^{-}$; E, B220$^{+}$ CD43$^{-}$ IgM$^{+}$ IgD$^{-}$; F, B220$^{+}$ CD43$^{-}$ IgM$^{+}$ IgD$^{+}$)

**D.** Absolute number of follicular (CD23$^{+}$ CD21$^{+}$) and marginal zone (CD23$^{-}$ CD21$^{high}$ B cells (B220$^{+}$ CD19$^{+}$) in the spleen (*n*= 5-8 mice per genotype, where each data point is a single mouse). Significance was determined by an unpaired two-tailed *t*-test (mean ± SEM). ns, not significant

**E.** Representative flow cytometry plots used to quantify B cell development in the bone marrow, gating on B220$^{+}$ CD43$^{+}$ for Hardy fractions A, B, C, and on B220$^{+}$ CD43$^{-}$ (Hardy fractions D, E, F).

**F.** Representative flow cytometry plots showing B220$^{+}$ CD19$^{+}$ cells used to quantify mature B cell populations in the spleen.

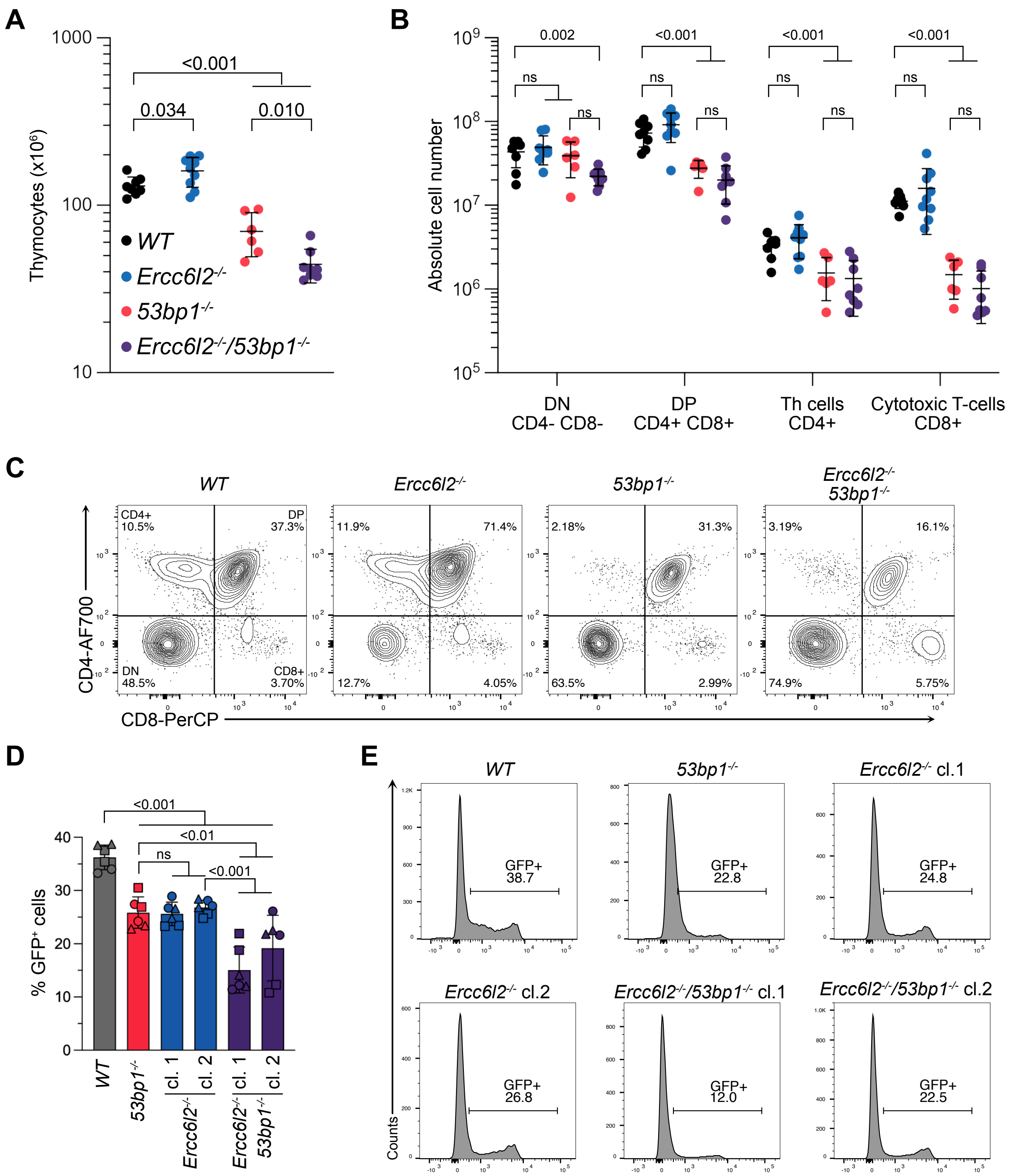


### Extended Data Fig 4

**Extended Data Fig. 4. T cell development in *53bp1$^{-/-}$ Ercc6l2$^{-/-}$* mice**

**A.** Absolute number of nucleated cells in the thymus (*n*= 6-8 mice per genotype, where each data point is a single mouse). Significance was determined by an unpaired two-tailed *t*-test (mean ± SEM).

**B.** Absolute number of thymic T cells (*n*= 6-8 mice per genotype, where each data point is a single mouse). Significance was determined by an unpaired two-tailed *t*-test (mean ± SEM). ns, not significant; DN, double-negative; DP, double-positive

**C.** Representative flow cytometry plots showing thymocytes used to quantify T cells in each developmental stage.

**D.** Flow cytometric analysis of GFP expression in *v-abl* pre-B cells of the indicated genotypes 96h after treatment with imatinib (*n*=3 biological replicates, with two technical replicates per experiment). Significance was determined by a one-way ANOVA with Tukey's correction (mean ± SD).

**E.** Representative flow cytometry plots showing quantification of GFP expression.

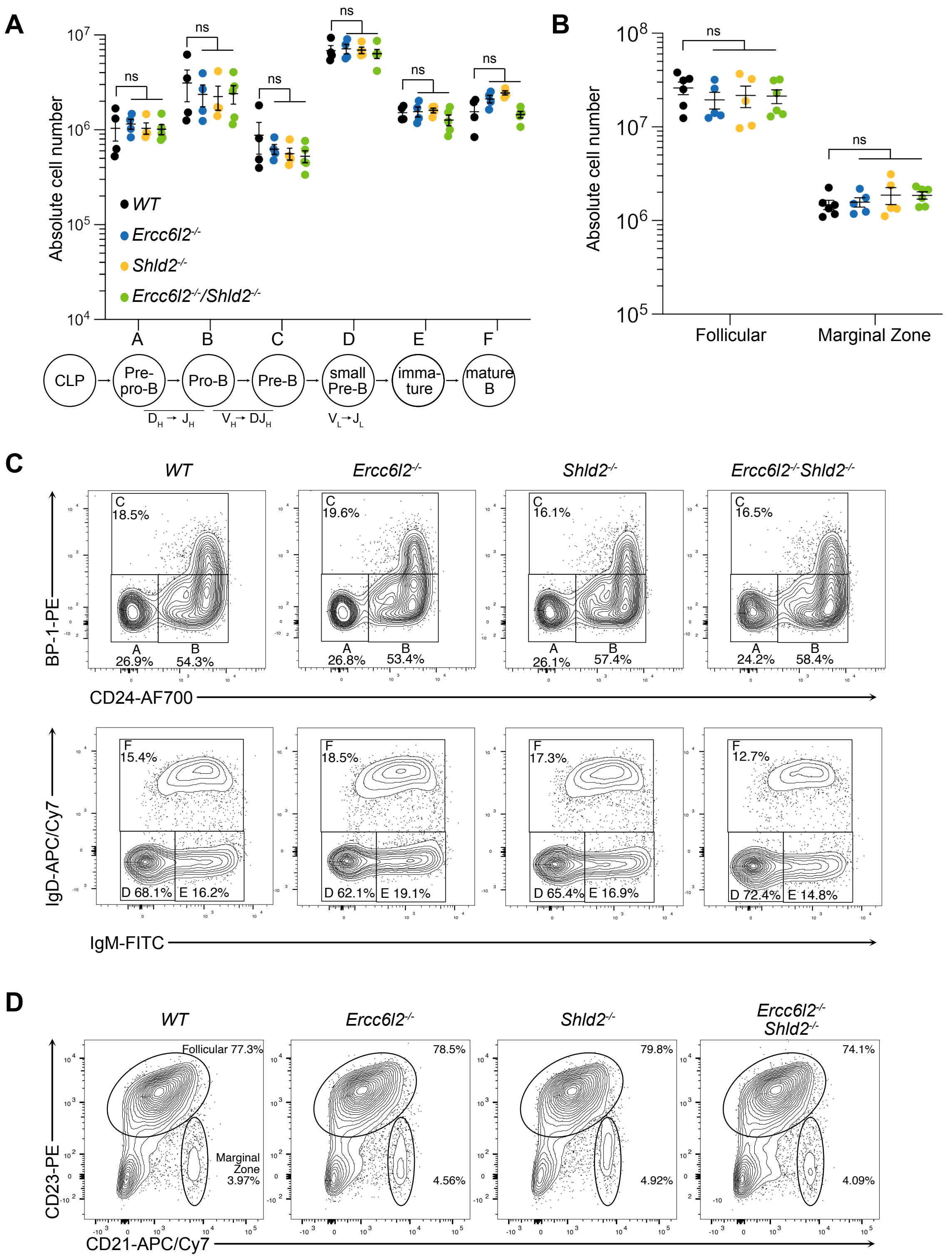


### Extended Data Fig 5

**Extended Data Fig. 5. B cell development in *Shld2$^{-/-}$ Ercc6l2$^{-/-}$* mice**

**A.** Absolute number of B220$^{+}$ B cell precursors in the bone marrow (one femur and one tibia) *n*= 5-8 mice per genotype, where each data point is a single mouse). Significance was determined by an unpaired two-tailed *t*-test (mean ± SEM). ns, not significant; A, B220$^{+}$CD43$^{+}$BP1$^{-}$CD24$^{-}$; B, B220$^{+}$ CD43$^{+}$ BP1$^{-}$CD24$^{+}$; C, B220$^{+}$ CD43$^{+}$ BP1$^{+}$ CD24$^{+}$; D, B220$^{+}$ CD43$^{-}$ IgM$^{-}$ IgD$^{-}$; E, B220$^{+}$ CD43$^{-}$ IgM$^{+}$ IgD$^{-}$; F, B220$^{+}$ CD43$^{-}$ IgM$^{+}$ IgD$^{+}$)

**B.** Absolute number of follicular (CD23$^{+}$ CD21$^{+}$) and marginal zone (CD23$^{-}$ CD21$^{high}$ B cells (B220$^{+}$ CD19$^{+}$) in the spleen (*n*= 5-8 mice per genotype, where each data point is a single mouse). Significance was determined by an unpaired two-tailed *t*-test (mean ± SEM). ns, not significant

**C.** Representative flow cytometry plots used to quantify B cell development in the bone marrow, gating on B220$^{+}$ CD43$^{+}$ for Hardy fractions A, B, C, and on B220$^{+}$ CD43$^{-}$ (Hardy fractions D, E, F).

**D.** Representative flow cytometry plots showing B220$^{+}$ CD19$^{+}$ cells used to quantify mature B cell populations in the spleen.

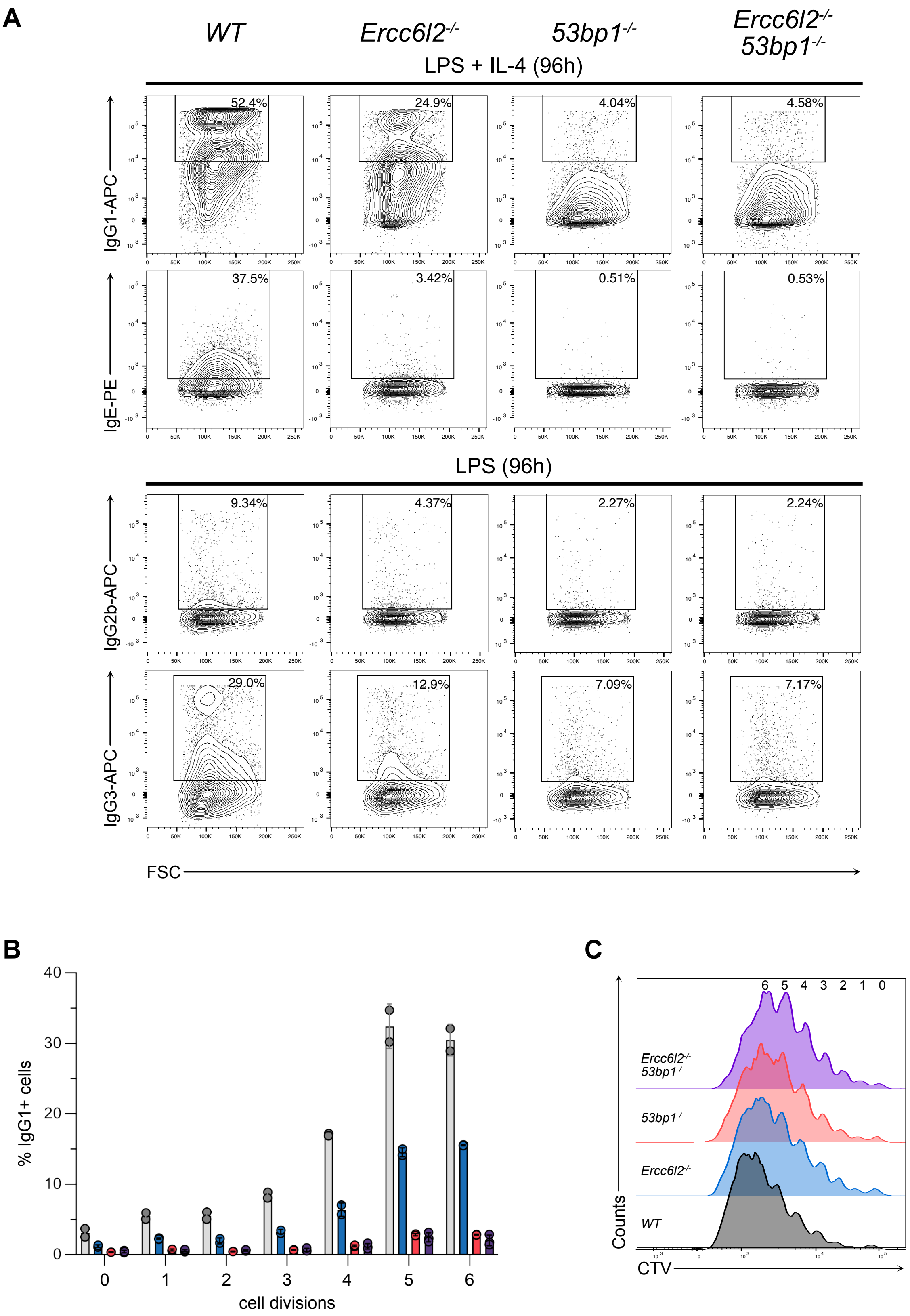


### Extended Data Fig 6

**Extended Data Fig. 6. ERCC6L2-53BP1 epistasis during CSR**

**A.** Representative flow cytometry showing purified splenic B cells stimulated with the indicated stimuli and stained for surface IgG1/G2b/G3 or IgE.

**B.** Percentage of $IgG1^+$ B cells in each cell division as determined by proliferation-associated CTV dilution (*n*= 2 mice per genotype, where each data point is a single mouse).

**C.** Representative flow cytometry plots of CTV dilution in purified splenic B cells stimulated with LPS and IL-4 for 96h.

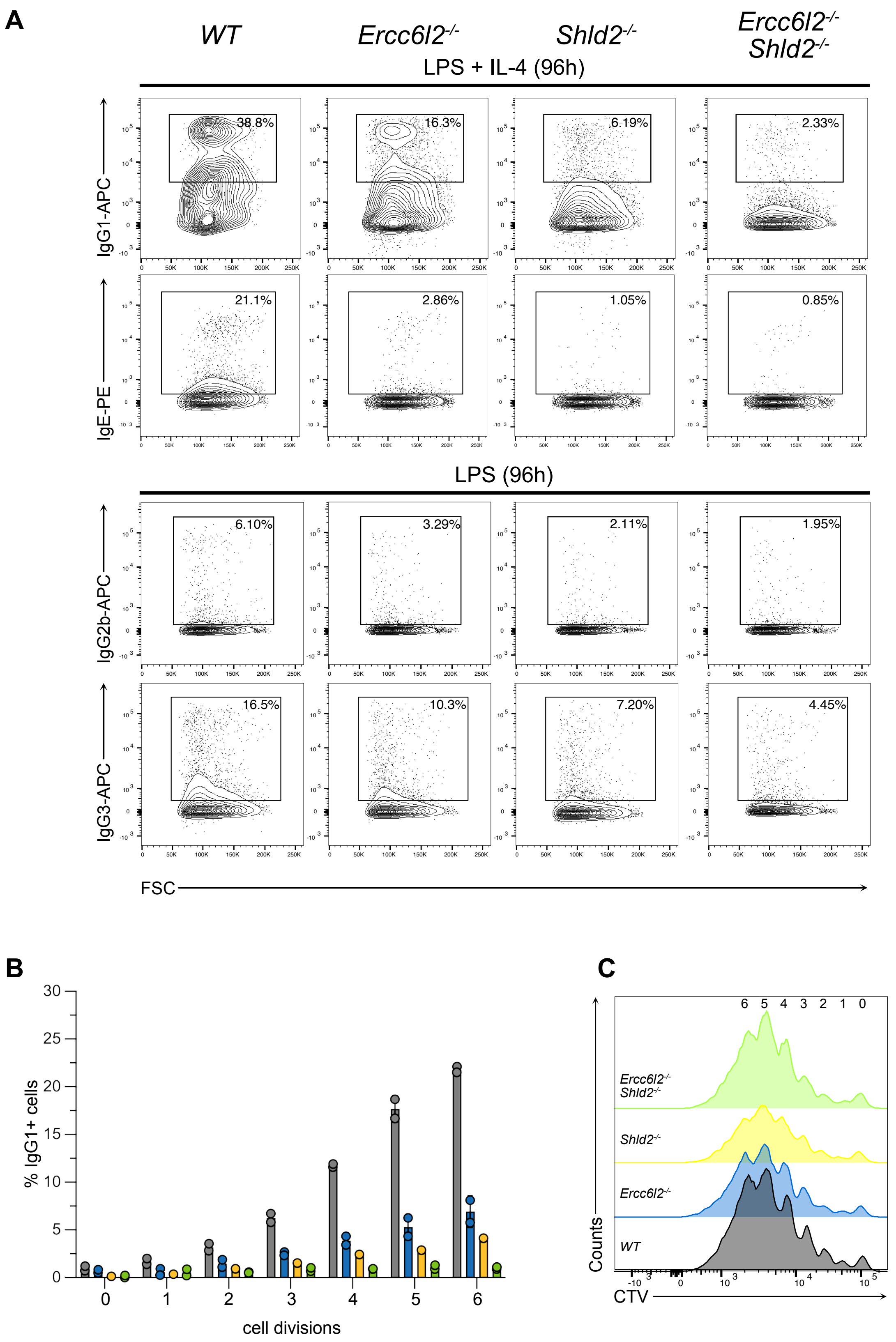


### Extended Data Fig 7

**Extended Data Fig. 7. ERCC6L2-shieldin synergy during CSR**
**A.** Representative flow cytometry showing purified splenic B cells stimulated with the indicated stimuli and stained for surface IgG1/G2b/G3 or IgE.
**B.** Percentage of $IgG1^+$ B cells in each cell division as determined by proliferation-associated CTV dilution (*n*= 2 mice per genotype, where each data point is a single mouse).
**C.** Representative flow cytometry plots of CTV dilution in purified splenic B cells stimulated with LPS and IL-4 for 96h.

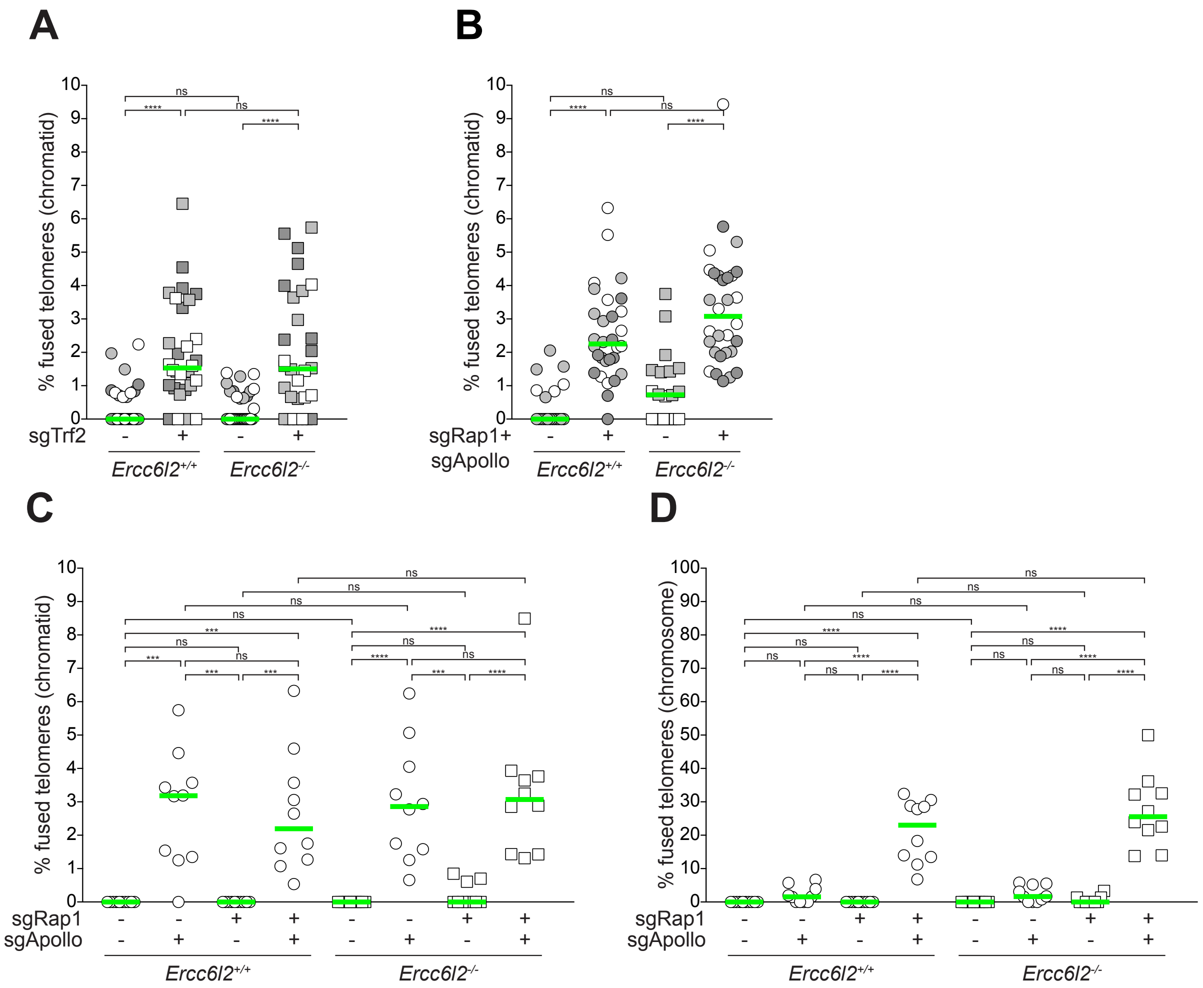


### Extended Data Fig 8

**Extended Data Fig. 8. ERCC6L2 stimulates c-NHEJ at staggered, but not blunt dysfunctional telomeres.**

**A.** Quantification of telomeres involved in chromatid fusions per metaphase in *Ercc6l2*$^{+/+}$ and *Ercc6l2*$^{-/-}$ MEFs 108 h after transduction with sgTrf2. Data from 3 independent experiments, 10 metaphases per experiment (n = 30 total), with median.

**B.** Quantification of telomeres involved in chromatid fusions per metaphase in *Ercc6l2*$^{+/+}$ and *Ercc6l2*$^{-/-}$ MEFs 120 h after transduction with sgRap1 and sgApollo. Data from 2 (no sgRNAs) or 3 (sgRAP1 + sgAPOLLO) independent experiments, 10 metaphases per experiment with median.

**C-D.** Quantification of telomeres involved in chromatid (c) or chromosome (d) fusions per metaphase in *Ercc6l2*$^{+/+}$ and *Ercc6l2*$^{-/-}$ MEFs 120 h after transduction with sgRap1 and/or sgApollo. Data from 1 representative experiment (n = 10 total), with median.

Ordinary one-way analysis of variance (ANOVA). *P ≤ 0.05, **P ≤ 0.01, ***P ≤ 0.001, ****P ≤ 0.0001; ns, not significant.

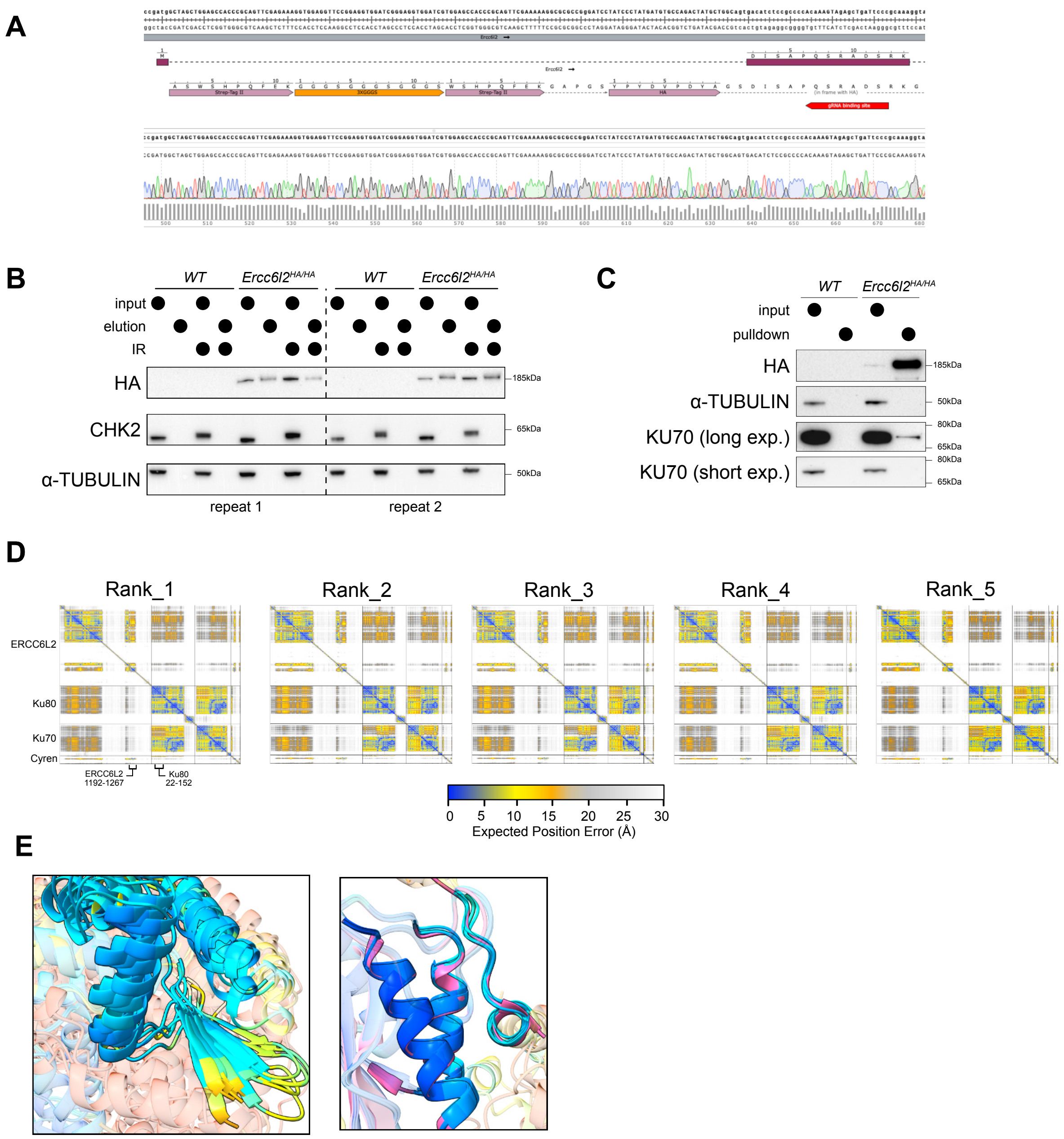


Extended Data Fig 9

**Extended Data Fig. 9. The ERCC6L2-MRI-KU complex supports classical NHEJ in CSR**

**A.** Top, schematic depicting the 5' coding region of the mouse *Ercc6l2* locus with bi-allelic targeted integrations of a TwinStrep-HA tagging cassette. Bottom, Sanger sequencing trace confirmation of the integrations in murine CH12-F3 cells.

**B.** Immunoblot analysis of CH12-F3 cell lysates from ERCC6L2 purification experiments used for subsequent analysis via LC-$MS^2$.

**C.** Immunoblot analysis of CH12-F3 cell lysates from an ERCC6L2 purification. Blot is representative of *n*=2 biological replicates.

**D.** PAE plots of all 5 predicted the ERCC6L2-MRI-Ku-DNA complexes showing expected position error. Peaks corresponding to MRI-ERCC6L2 and MRI interface are highlighted in red box.

**E.** Superimposed ERCC6L2-MRI binding site of all AF2 structures (left) and Ku80-MRI binding sites (right).
